# Abrogating TGFβ signaling in TCR-engineered T cells and enhancing antigen processing by tumor cells promotes sustained therapeutic activity in pancreatic ductal adenocarcinoma

**DOI:** 10.1101/2025.06.04.657887

**Authors:** Alexander K. Tsai, Meagan R. Rollins, Madeline A. Ellefson, Zoe C. Schmiechen, Adam L. Burrack, Ayaka Hulbert, Guhan Qian, Hongrong Zhang, Paolo P. Provenzano, Eduardo Cruz- Hinojoza, Jonah Z. Butler, Olivia C. Ghirardelli Smith, Stephen D. O’Flanagan, Jenny Krause, Grant H. Hickok, David Masopust, Sunil R. Hingorani, Philip D. Greenberg, Ingunn M. Stromnes

## Abstract

Pancreatic ductal adenocarcinoma (PDA) is a deadly malignancy with limited effective therapies. Adoptive cell therapy (ACT) is a promising treatment modality for patients with solid tumors but has been limited by the highly fibroinflammatory and immunosuppressive tumor microenvironment (TME). Transforming growth factor-β (TGFβ) participates in the inordinately suppressive TME in PDA. Here, we test the impact of selective *Tgfbr2* deletion using CRISPR/Cas9 or genetic approaches in mesothelin (Msln)-specific T cell receptor (TCR) engineered T cells during ACT of PDA. Abrogating TGFβ signaling augmented TCR-engineered T cell accumulation in autochthonous and orthotopic PDA models and promoted terminal effector T cells, although this largely required inclusion of a vaccine at the time of T cell transfer. While loss of *Tgfbr2* impaired CD103 upregulation, it only modestly impaired donor T cell central, tissue resident, or Tcf1^+^Slamf6^+^ stem-like memory T cell formation. These attributes ultimately result in heightened functional capacity and delayed tumor growth. Unexpectedly, however, most tumor-infiltrating engineered T cells failed to differentiate into PD-1^+^Lag3^+^ exhausted T cells (T_EX_) regardless of TGFβR2 expression and despite abundant Msln protein expression by PDA cells. Forcing Msln epitope processing in *KPC* tumor cells promoted donor T cell accumulation, acquisition of PD-1 and Lag3, increased IFNγ production by TCR-engineered T cells refractory to TGFβ and bypassed the vaccine requirement for therapeutic efficacy. Thus, promoting increased antigen processing/presentation by tumor cells while abrogating *Tgfbr2* in engineered T cells can sustain donor T cell function in the suppressive TME and enhance the therapeutic efficacy of ACT. Our study supports pursuit of strategies that modulate tumor intrinsic antigen processing while relieving T cell suppression to safely promote the antitumor activity of TCR-engineered T cells.

## INTRODUCTION

Pancreatic ductal adenocarcinoma (PDA) is the third deadliest malignancy in the U.S. (1). Mortality rates are predicted to rise in the coming years, in part due to increasing trends in PDA incidence over the past two decades (1). Most patients are diagnosed with unresectable and/or metastatic PDA and have a median overall survival of less than one year (2–4). Though new chemotherapy regimens and targeted therapies have been developed (4–6), they provide only incremental advantages over prior standards (2, 3). Thus, there remains an unmet and urgent need to develop more effective therapies.

Immune checkpoint blockade (ICB) therapies are approved for the treatment of many malignancies but lack efficacy in PDA (7–9). Failure of ICB efficacy in PDA has largely been attributed to the profoundly suppressive and fibroinflammatory tumor microenvironment (TME), which impairs tumor-infiltrating lymphocyte (TIL) infiltration, activation, and effector function (10). Moreover, PDA exhibits relatively low tumor mutational burden, resulting in a paucity of mutation-derived neoantigens (11–13). Adoptive cell therapy (ACT) in which T cells are genetically modified to express a tumor-reactive antigen receptor is a provocative approach for PDA. While chimeric antigen receptor T cell (CAR-T) therapies have improved outcomes in various hematologic malignancies (14), application of CAR-T cell therapy in solid tumors, including PDA, has had limited success (15). T cell receptor (TCR) engineered T cell therapy has shown some promise in both PDA (16) and other GI malignancies (17, 18). Though somewhat confined by the requirement for HLA allele matching, this approach allows for targeting of intracellular antigens and trials using MAGE-A4 targeting TCRs have shown efficacy leading to approval for treatment of synovial sarcoma (19). Additional trials examining various antigens using TCR-engineered ACT or through enrichment of neoantigen-specific TILs are ongoing (18, 20, 21). However, limited numbers of patients have been treated thus far, and there remains a critical knowledge gap as to how to safely improve ACT for PDA patient treatment, presumably because of the myriad of obstacles presented by the suppressive TME.

We previously employed an autochthonous genetically engineered *Kras*^G12D/+^;*Trp53*^R172H/+^;*p48*-Cre (*KPC*) mouse model of PDA that recapitulates hallmark features of the human disease (22) to evaluate ACT efficacy. Semi-monthly infusions of CD8^+^ T cells retrovirally transduced to express a high-affinity murine mesothelin (Msln)_406-414_:H2D^b^-specific TCR (clone 1045) combined with lymphodepletion and an irradiated peptide-pulsed cell vaccine resulted in intratumoral donor T cell accumulation, objective responses, and a near-doubling of *KPC* survival (23). However, intratumoral TCR-engineered T cell number decreased between infusions and persisting donor T cells were rendered dysfunctional, or exhausted (23). Further, donor T cell function was neither rescued by immune checkpoint blockade combinations (24) nor through depletion or activation of tumor-associated macrophages (25).

Transforming growth factor-β (TGFβ) is a potent immunosuppressive cytokine that is elevated in PDA. TGFβ promotes epithelial-mesenchymal transition (EMT) and activates pro-tumor cancer associated fibroblasts (CAFs) (26–28). Paradoxically, while TGFβ can promote malignant hallmarks, loss of TGFβ can also cooperate with other alterations to promote oncogenesis. Indeed, *SMAD4*, an intracellular messenger downstream of TGFβ signaling, is mutated or lost in over half of PDA cases and contributes to early PDA carcinogenesis (29, 30). As such, systemic inhibition of TGFβ may have undesirable pro-tumor effects in some scenarios. As TGFβ is essential in normal biology and homeostasis of all organ systems, systemic inhibition can also result in adverse effects, particularly cardiac toxicity (31, 32). One approach to circumventing the toxicities observed with systemic TGFβ inhibition is to use targeted strategies, ideally with specificity for individual cell subsets.

This approach is ideal for ACT, where T cells can be engineered and modified *ex vivo* prior to reinfusion into patients. In CD8^+^ T cells, TGFβ limits T cell activation partially through Src homology region 2 domain-containing phosphatase-1 (SHP-1) upregulation which blunts T cell receptor (TCR) signaling (33, 34). This, along with other mechanisms reduces CD8^+^ T cell proliferation, effector T cell differentiation (35, 36), and impairs effector function including production of interferon-γ (IFNγ), interleukin-2 (IL-2), and granzyme B (GzmB) (37–39).

As TGFβ is abundant in human PDA and potently immunosuppressive (26), we hypothesized that it may pose a hierarchical barrier to ACT therapeutic activity. Here, we extend our prior findings of ACT targeting Msln in PDA by elucidating the impact of abrogating TGFβ signaling specifically in adoptively transferred T cells using PDA animal models. We find T cell intrinsic disruption of TGFβR2 promotes TCR-engineered T cell differentiation toward effector T cells with improved antitumor function. However, we also identify that enhancing Msln epitope processing in tumor cells is critical to improve the therapeutic efficacy of such TCR-engineered T cells.

## METHODS

### Animals

The University of Minnesota Institutional Animal Care and Use Committee (IACUC) approved all animal studies. Animals were maintained in specific pathogen-free conditions and experiments conformed to ethical regulations for animal testing and research. 6-to 8-week-old male and female C57BL/6J (000664), P14 (037394-JAX), Tgfbr2^Flox/Flox^ (012603), and *dLck*-Cre (012837) mice were purchased from The Jackson Laboratory. Xcr1-Venus mice were kindly provided by Dr. Brian Fife (University of Minnesota). 1045 T cell receptor exchange (TRex) mice that express the high affinity Msln_406-414_:H-2D^b^-specific TCR within the physiological *Trac* locus crossed to Thy1.1 were described (40). Previously described *Kras*^LSL-G12D/+^;*Trp53*^LSL-R172H/+^;*p48*-Cre (*KPC*) mice speed-backcrossed to the C57Bl6/J strain (23) were maintained in facilities at the University of Minnesota. Mice were bred to achieve desired genotypes, which were determined using custom probes and sequencing provided by Transnetyx. Animals were maintained in SPF conditions at the University of Minnesota Research Animals Resources facility with continual access to food and water and kept on a 12-hour light-dark cycle.

### PDA tumor cell culture

*KPC*2 tumor cells were established from a primary tumor derived from a female *KPC* mouse as previously described (41). Additional cell lines, including *KPC*271 and *KPC*451, were similarly derived from independent *KPC* mice. *KPC*2 cells were transduced with Click Beetle Red (CBR) luciferase to generate the *KPC*2a cell line (41). For two-photon imaging, *KPC*2 cells were transduced with mScarlet cloned in the MIGR1 plasmid (kindly provided by Dr. Marc Jenkins, University of Minnesota). Transduced cells were sorted using a FACS Aria II (BD) to >95% purity. For cytotoxicity assays, *KPC*2 cells were transduced with Incucyte Nuclight NIR lentivirus (Sartorious) and subsequent cells were maintained under antibiotic selection with media supplemented with 1 μg/mL puromycin (Gibco). The *KPC*2^Msln-ER^ cell line was generated by transducing *KPC*2 cells to express Msln_406-414_ using a previously described “PresentER” plasmid (42). *KPC*2^Msln-ER^ cells were sorted to >95% purity using a MACSQuant Tyto cell sorter (Miltenyi Biotec). All cell lines were maintained at 37°C and 5% CO_2_ below passage 20 in pancreatic tumor media (PTM): DMEM-F12 (Gibco), 10% fetal bovine serum (FBS, Gibco), 2.5 mg/mL amphotericin B (Gibco), 100 U/mL penicillin/streptomycin (Gibco), 2.5 g dextrose (Thermo Fisher).

### In vitro proliferation assay

Proliferation was assessed as described previously (43). Splenocytes were isolated and washed twice with PBS. Cells were stained with 1 μM CellTrace Violet (CTV, Thermo Fisher) in PBS for 20 minutes at 37°C and subsequently quenched with 4 volumes of 4°C T cell media (TCM): DMEM (Gibco), 10% FBS, 4 mM L-glutamine (Gibco), 1x non-essential amino acids (NEAA, Gibco), 50 μM 2-mercaptoethanol (Sigma Aldrich), 100 U/mL penicillin-streptomycin (Life Technologies). CTV-labeled splenocytes were stimulated with titrating concentrations of Msln_406-414_ peptide (Genscript) and 10 nM recombinant human interleukin-2 (rhIL-2, Peprotech) for 72 hours.

### In vitro cytokine production assay

Splenocytes were isolated and primed with 5 μg/mL of Msln_406-414_ and 10 nM rhIL-2 for 72 hours. Subsequently, cells were restimulated with 1 μg/mL plate-bound anti-CD3 (Clone 17A2, BD) ± 20 ng/mL Tgfβ1 (R&D Systems) for 24 hours. 1:1000 dilution of GolgiPlug (BD) and 1:1500 dilution of GolgiStop (BD) was added for the final 4.5 hours of restimulation.

### Cytotoxicity assay

Splenocytes were primed with 5 μg/mL of Msln_406-414_ for 72 hours. *KPC*2 NIR^+^ were plated and allowed to adhere for 6 hours. CD8^+^ T cells were enriched using negative selection (Miltenyi) and subsequently added to previously plated *KPC*2 tumor cells at a 5:1 Effector:Target (E:T) ratio. Tumor cell growth was monitored using an Incucyte SX5 instrument (Sartorius) for 72 hours.

### Co-culture assay

Splenocytes were primed with 1 μg/mL of Msln_406-414_ for 48 hours. Primed cells were washed, and media was replaced with TCM + 10 nM rhIL-2 for 24 hours. Tumor cells were plated ± 100 μg/mL IFNγ (R&D Systems) for 24 hours. For some conditions, tumor cells were pulsed with 5 μg/mL of Msln_406-414_ for 1 hour and subsequently washed with PTM. Primed and rested T cells were added to tumor cells at a 5:1 E:T ratio and co-cultured for 24 hours. 1:1000 dilution of GolgiPlug and 1:1500 dilution of GolgiStop was added for the final 4.5 hours of co-culture.

### Orthotopic tumor implantation

Orthotopic *KPC*2 tumor implantation was performed as previously described (41, 44–46). Briefly, 1 × 10^5^ tumor cells in 20 μL of 60% Matrigel (Corning) were injected into the pancreas via insulin syringe (Covidien) following surgical peritoneal access in anesthetized mice with 2-5% isoflurane and pretreated with 1 mg/kg buprenorphine for analgesia according to approved protocols. Following tumor implantation, the peritoneum was closed with 6-0 sutures (Ethicon), and the overlying skin was closed using 9-mm wound clips (Fine Science Tools). Mice were treated with 10 mg/kg carprofen immediately following surgery and at 24 and 48 hours after tumor implantation and monitored for 5 days to ensure adequate wound healing.

### Adoptive cell therapy protocols

For experiments in the genetically engineered *KPC* mouse model in Figure 1, congenic P14 *dLck*-Cre *x Tgfbr2*^WT/WT^ or P14 *dLck*-Cre x *Tgfbr2*^Flox/Flox^ TCR transgenic T cells were retrovirally transduced with 1045 TCR and subsequently expanded as previously described (23). Prior to ACT, 180 mg/kg cyclophosphamide (Amneal Pharmaceuticals) was delivered intraperitoneally (i.p) for lymphodepletion. A 1:1 mixture of Tgfbr2 WT and KO (2.5 × 10^6^ of each type of engineered T cells) was delivered i.p along with 1 × 10^7^ irradiated antigen presenting cells (APC) pulsed with Msln_406-414_ peptide (23). For all other ACT protocols, mice received 120 mg/kg cyclophosphamide i.p followed by 5 × 10^6^ Thy1.1^+^*dLck*-Cre x *Tgfbr2*^WT/WT^ wild type (WT) or *dLck*-Cre x *Tgfbr2*^Flox/Flox^ CD8^+^ T cells isolated from 1045 TRex mice and primed with 1-5 μg/mL of Msln_406-414_ 72 hours prior to ACT. Where indicated, mice also received either BiVax [100 μg Msln_406-414_ + 50 mg Poly I:C (Tocris)] (47) or TriVax [100 μg Msln_406-414_ + 50 mg Poly I:C + 50 μg anti-CD40 (Clone FGK4.5, BioXcell)] (48). All mice also received 5 μg rhIL-2 with T cell delivery in addition to 48 and 96 hours after ACT.

**Figure 1.**
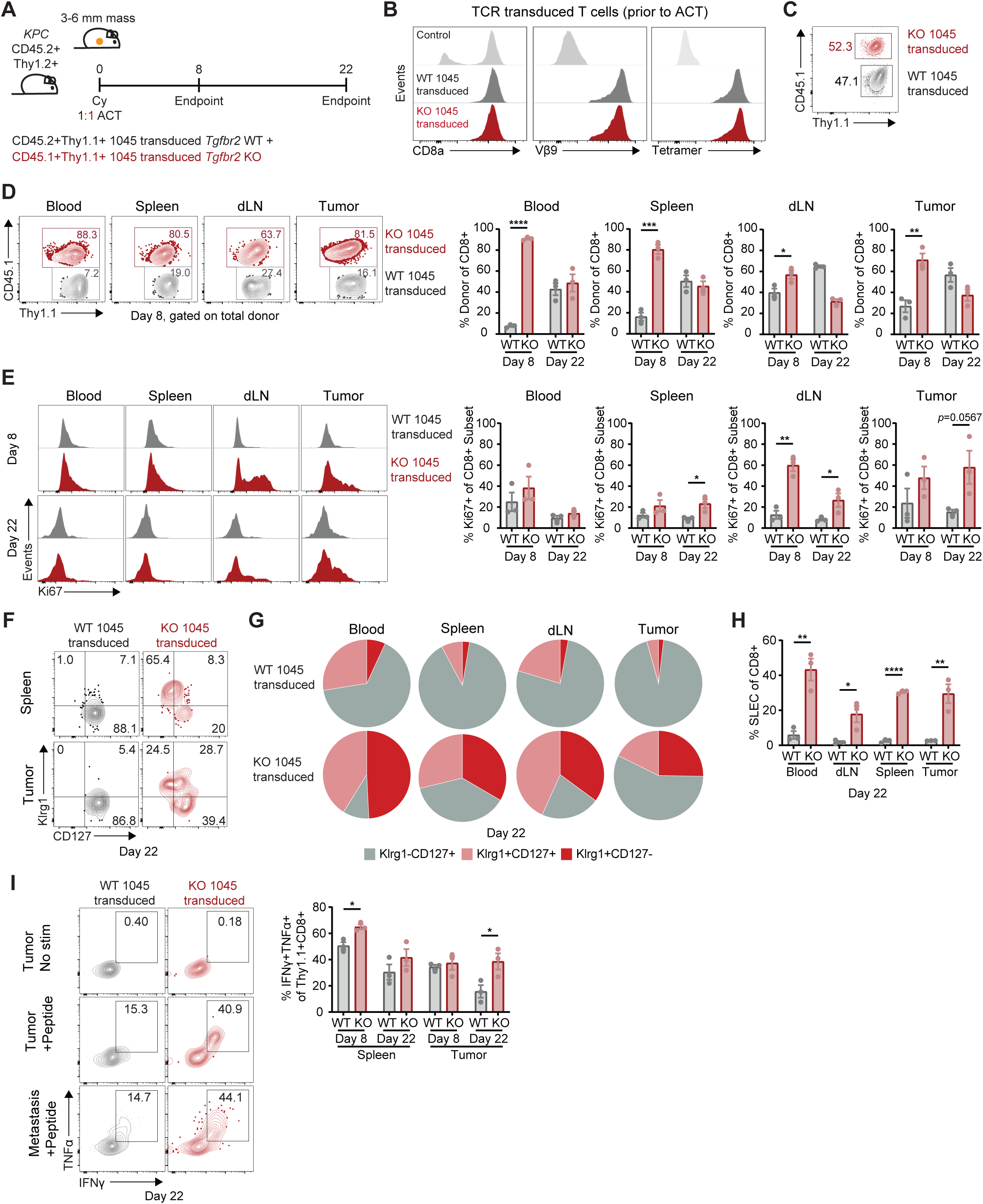
TGFβ impairs TCR-engineered T cell expansion, proliferation, and effector function in autochthonous PDA. **A)** Schematic illustrating ACT using co-transfer of 1045 transduced *Tgfbr2* WT and KO T cells (2.5 × 10^6^ of each cell type) and vaccination with Msln_406-414_-pulsed APCs (1 × 10^7^ cells) in *KPC* mice harboring 3-6 mm tumors following lymphodepletion with cyclophosphamide (180 mg/kg). **B)** 1045 TCR expression assessed with Vβ9 staining (middle) and Msln_406-414_:H2D^b^ tetramer binding (right) in transduced *Tgfbr2* WT and KO cells compared to untransduced cells. **C)** Frequency of CD45.2^+^Thy1.1^+^ *Tgfbr2* WT cells and CD45.1^+^Thy1.1^+^ *Tgfbr2* KO cells prior to ACT. **D)** Representative plots (day 8, left) depicting 1045 transduced *Tgfbr2* WT and KO cells of total donor cells in blood, spleen, dLN, and tumor on days 8 and 22 (right). **E)** Representative histograms (left) depicting Ki67^+^ frequency of 1045 transduced *Tgfbr2* WT or KO donor cells (right). **F-G)** Representative plots (F) showing Klrg1^-^CD127^+^, Klrg1^+^CD127^+^, or Klrg1^+^CD127^-^ mean frequency of 1045 transduced *Tgfbr2* WT or KO donor cells on day 22 (G). **H)** SLEC (Klrg1^+^CD127^-^) frequency of 1045 transduced *Tgfbr2* WT or KO donor cells on day 22. **I)** Representative plots (left, including a liver metastasis) depicting IFNγ^+^TNFα^+^ frequency of 1045 transduced *Tgfbr2* WT or KO donor cells following restimulation with Msln_406-414_ on day 22. Dots are individual animals and n=3/group in all panels with mean ± SEM shown. Student’s t-test was used for comparisons in all panels.

### CRISPR/Cas9 mediated deletion of Tgfbr2

Two custom guide RNAs (gRNA), 5’-G*A*C*CGCACCGCCAUUGU and 5’-C*U*C*AGUUAACAGUGAUG (Synthego), were designed targeting exon 1 of *Tgfbr2*. gRNAs were suspended at 100 pmol/μL and complexed at a 3:1 ratio with TrueCut Cas9 V2 (Invitrogen). 1045 TRex T cells previously primed with 10 μg/mL Msln_406-414_ peptide 72 hours prior were suspended in RT P4 nucleofection buffer (Lonza), 4 μM Alt-R Cas9 electroporation enhancer (Lonza), and complexed gRNA with Cas9 in nucleofection cuvettes (Lonza). Electroporation was performed using a 4D Nucleofector (Lonza) using program CM137. To analyze editing efficiency, DNA was isolated from edited cells using DNeasy Blood and Tissue Kit (Qiagen) according to the manufacturer’s instructions. Sanger sequencing (Eurofins) was performed with primer 5’-TGTCGCGGCTGCATATC and editing efficiency was determined using Inference of CRISPR Edits (ICE) online software (Synthego). Edited cells were adoptively transferred within 24 hours of electroporation.

### Two-photon imaging and analysis

*KPC*2 mScarlet^+^ tumors were implanted in Xcr1-Venus mice and subsequently isolated following treatment with ACT plus TriVax as described above. The entire tumor was embedded in 0.5% agarose. Embedded tumors were cut at 3 distinct depths into 20-micron slices using a microtome, mounted on a coverslip, and equilibrated in warm oxygenated RPMI 1640 containing 5% FBS for 30 minutes. Movies were acquired using a MP SP5 two photon microscope TCS (Leica) equipped with a Mai Tai HP Deep See lasers (SpectraPhysics), an 8,000-Hz resonant scanner, a 25x 3 /0.95 NA objective, and two non-descanned and two hybrid detectors.

During imaging, continuous oxygenated DMEM high glucose media lacking phenol red (Hyclone) was exchanged in the chamber containing the sample. Tissue was excited at 860 or 890 nm and multiple fluorophores were imaged using custom dichroic mirrors with the following collections: SHG < 440nm, mScarlet 569-593 nm, GFP 500-520 nm, Venus 515-528 nm. For each individual tumor, videos were collected over a 30 minute period in 3-5 different geographic locations separated by at least 1 mm for a total of sixty frames. Within each of these macroscopic locations, 2-3 positions were imaged concurrently.

For two-photon analysis, DC and T cells were separately tracked using the automatic tracking algorithm TrackMate in FIJI. Individual cells were first identified using the Laplacian of Gaussian (LoG) detector and cell movement was then tracked using the Linear Assignment Problem (LAP) tracker. Cell motility was calculated by fitting the mean square displacement (MSD) from the track data with the persistent random walk model (PRWM), which can be described as follows, where *S* is the migration speed and *P* is the persistence time:

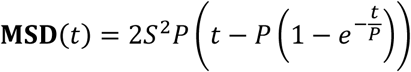

The motility coefficient is given as follows, where *n_d_* is of unity dimension:

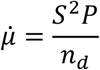

Thus, the total motility coefficient is the sum of motility components and is given as:

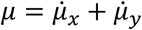

Statistical outliers in motility coefficients were removed by ROUT (Q=1%) in GraphPad Prism software (version 10). Individual cell track features like displacement, total distance traveled, confinement ratio, mean track speed, and mean directional change rate were used to further characterize cell movement.

### Intravenous labeling and single cell suspension generation

IV labeling was performed as described previously (46, 49). Briefly, 3 μg of anti-CD45 antibody was injected intravenously in anesthetized mice 3 minutes prior to euthanasia. In all mice, single cell suspensions were generated as previously described (46). Lymph nodes (LN) were mechanically dissociated over 40 μM filters. Tumors and pancreases were minced and incubated with 0.5 mg/mL Collagenase IV (Gibco) in DMEM for 15 minutes in an Incu-Shaker (Carolina) at 37°C. Digested tumor and pancreas fragments were subsequently mechanically dissociated over 70 μM filters. Spleens were mechanically dissociated followed by RBC lysis using Tris-ammonium chloride (ACK) lysis buffer (Gibco) and subsequent quenching with TCM. Mechanically dissociated livers and minced lungs were subjected to 1 hour of enzymatic digestion using Collagenase type I (Worthington Biochemicals) and subsequently dissociated using a gentleMACS tissue dissociator (Miltenyi). Tissues were passed over 70 μM filters and mononuclear cells were enriched via Percoll gradient.

Intraepithelial lymphocytes (IEL) within small intestines (SI) were isolated using a method established in Dr. David Masopust’s laboratory (50). Briefly, SI peyer’s patches and contents were removed followed by longitudinal dissection to expose the inner mucosal surface. Cleaned SI were segmented and subsequently incubated for 30 minutes with stirring at 37°C with 0.154 mg/ml dithioerythritol (DTE, Sigma Aldrich) in 10% Hank’s Balanced Salt Solution [HBSS, without sodium bicarbonate, calcium, and magnesium from (Corning)]. IEL were released by vortexing at high-speed and the supernatant was collected and enriched on a Percoll gradient. All single cell suspensions were washed twice with TCM prior to further processing.

### Ex vivo cytokine production

Splenic and intratumoral T cells were incubated at 37°C / 5% CO_2_ for 4.5 hours ± 1-5 μg/mL of Msln_406-414_ or 1:500 dilution of cell stimulation cocktail (CSC, Tonbo). Cells were cultured with 1:1000 dilution of GolgiPlug and 1:1500 dilution of GolgiStop.

### Flow cytometric analyses

Antibodies used for flow cytometry are listed in **Supplemental Table 1**. CB_101-109_:H2D^b^-BV421 tetramer was generated as previously described (41). Cultured cells or single cell suspensions were washed twice in FACS buffer [phosphate buffer solution (PBS, Gibco), 2% FBS, 0.01% sodium azide (Ricca Chemical Company)].

Cells were then incubated with a mixture of antibodies diluted at 1:100 or 1:200 in FACS buffer for 1 hour at room temperature (single cell suspensions) or 45 minutes at 4°C (cultured cells) protected from light. Fixation and permeabilization was performed for conditions requiring intracellular staining of cytokines using a Cytofix/Cytoperm fixation/permeabilization kit (BD). For intracellular staining of transcription factors, a Foxp3 fixation/permeabilization kit (Tonbo) was used. Antibodies for intracellular staining were diluted at 1:100 to 1:300 in perm/wash buffer and incubated overnight at 4°C protected from light. Following staining, all cells were washed twice and resuspended in FACS buffer and analyzed using a 5-laser Cytek Aurora instrument. Data acquisition was performed using SpectroFlo software (Cytek) and analysis was performed using FlowJo software (Version 10).

### qPCR

*KPC* tumor cell lines were isolated from culture and RNA was extracted with QIAwave RNA Mini Kits (Qiagen). Total RNA concentration was determined from A260/A280 values obtained using a NanoDrop instrument (Thermo Fisher). cDNA was synthesized from RNA using High-Capacity RNA-to-cDNA Reverse Transcription Kits (Applied Biosystems). All qPCR reactions were performed in triplicate. Each reaction consisted of 500 nM forward and reverse primers (listed in Supplemental Table 1), 10 ng cDNA, PowerTrack SYBR Green Master Mix (Applied Biosystems), and nuclease-free water. qPCR amplification was performed in a C1000 Touch thermal cycler (Bio-Rad) programmed with the following settings: 1 cycle of enzyme activation at 95°C for 2 minutes; 40 cycles of denaturation at 95°C for 15 seconds and annealing at 60°C for 1 minute; and 1 melt curve cycle with a temperature range of 60-95°C at 0.5°C increments for 15 seconds. Fold change calculations were performed using the 2-ΔΔCt method, in which Gapdh served as a housekeeping transcript.

### Immunofluorescent microscopy

Tissue samples were preserved in OCT, rapidly frozen, and stored at -80°C. 7 μM tissue sections were prepared using a cryostat (Leica), mounted on glass slides, and fixed with 100% acetone for 10 minutes. Prepared slides were warmed to room temperature and blocked with 5% bovine serum albumin (BSA) in PBS for 1 hour at room temperature in a humidified chamber. Blocked samples were incubated with primary antibodies (listed in Supplemental Table 1) at 1:100 or 1:200 dilutions in 1% BSA for 1 hour at room temperature. ProLong Gold antifade with DAPI (Invitrogen) was applied and samples were stored at 4°C prior to imaging on a Leica DM4 B upright microscope.

### High-resolution ultrasound

Mice were anesthetized using a continuous flow of 2% to 4% isoflurane, and abdominal hair was removed using Nair (Church & Dwight). Tumors were identified based on anatomic landmarks, hypoechoic features, and clear tumor boundaries using high-resolution ultrasound (Vevo F2, VisualSonics) (46, 51). Tumor volume was calculated using a modified ellipsoidal formula: 0.5(length × width^2^). Euthanasia was performed when tumors reached ≥500 mm^3^ in volume.

### Statistical analyses

Unpaired two-tailed Student’s t-tests were used when comparing two groups. For two-photon migration evaluations, Welch’s t-test was used in some cases based on population variance. One-way ANOVA with Tukey posttests were used when comparing more than two groups. All measurements were performed independently with statistical significance groups of \**p*<0.05, \*\**p*<0.01, \*\*\**p*<0.001, and \*\*\*\**p*<0.0001 using GraphPad Prism software (version 10).

## RESULTS

### TGFβ impairs TCR-engineered T cell expansion, proliferation, and effector function in autochthonous PDA

To abrogate TGFβ signaling in tumor-specific lymphocytes, CD8^+^ T cells isolated from congenically distinct CD45.1^-^Thy1.1^+^ wild type (WT) or CD45.1^+^Thy1.1^+^ *Tgfbr2*^Flox/Flox^ *x dLck*-Cre (KO) P14 mice were retrovirally transduced with the 1045 TCR, expanded *in vitro*, and co-transferred at a 1:1 ratio into *KPC* mice. Hosts with ultrasound-defined 3-6 mm pancreatic tumor masses were treated with cyclophosphamide 6 hours prior to T cells, which were delivered with a cell-based vaccine consisting of Msln_406-414_ peptide-pulsed irradiated splenocytes (23) (**Figure 1A-C**). On day 8, 1045 KO donor T cells significantly outcompeted WT counterparts in circulation, spleen, draining lymph node (dLN), and tumor (**Figure 1D**). By day 22, however, frequencies of WT and KO T cells were similar in all tissues, despite the loss of *Tgfbr2* increasing donor T cell proliferation in all tissues as assessed by Ki67 (**Figure 1E**). Klrg1 and CD127 (IL-7Rα) were used to delineate donor memory precursor effector cells (MPEC) from short-lived effector cells (SLEC) (52) on day 22 (**Figure 1F-H**). A higher frequency of 1045 KO T cells displayed a SLEC phenotype or co-expressed Klrg1 and CD127 – a potential transitory state – as compared to WT T cells. Specifically, on day 22, SLEC comprised nearly 30% of KO donor T cells, which was nearly 10-fold greater than detected in WT donor T cells. To test if TGFβ impacted T cell function, donor T cells were stimulated with Msln_406-414_ peptide *ex vivo* and cytokine production was measured (**Figure 1I**). On day 22, a 2-fold higher frequency of 1045 KO T cells infiltrating autochthonous tumors and a liver metastasis produced effector cytokines as compared to 1045 WT T cells. Together, deletion of *Tgfbr2* in TCR-engineered CD8^+^ T cells improved donor T cell accumulation, proliferation, and effector function in autochthonous PDA.

### Abrogating T cell-intrinsic TGFβ improves endogenous neoantigen-specific T cells in orthotopic PDA

Accurately assessing ACT efficacy in autochthonous PDA is challenging owing to the spontaneous and heterogeneous nature of developing tumors. Thus, we evaluated the suppressive impact of TGFβ by orthotopic implantation of our previously described *KPC*2a PDA cell line (41) in *Tgfbr2*^Flox/Flox^ *dLck*-Cre^-^ (*Tgfbr2* WT) or *Tgfbr2*^Flox/Flox^ *dLck*-Cre^+^ (*Tgfbr2* KO) hosts. Use of *Tgfbr2*^Flox/Flox^ *dLck*-Cre mice allowed for the specific abrogation of TGFβ exclusively within T cells. *KPC*2a expresses click beetle red luciferase (CBR) which serves as a model neoantigen and results in development of endogenous CD8^+^ T cells specific for CB_101-109_:H2D^b^ that can be tracked using tetramer (41). *KPC*2a tumors were allowed to establish and progress for 14 days prior to evaluation of tumor-specific T cells (**Figure 2A**). No detectable tumors were recovered from 3 of 9 *Tgfbr2* KO mice, and tumors were significantly decreased in size in *Tgfbr2* KO compared to *Tgfbr2* WT animals (**Figure 2B**). CD8^+^ T cells were examined in non-draining inguinal lymph nodes (ndLN), dLN, spleens, and tumors (**Figure 2C**). Reduced CD8^+^ T cell frequencies were observed in ndLN and dLN of *Tgfbr2* KO mice, whereas CD8^+^ T cells were slightly enriched in spleens of *Tgfbr2* KO mice and intratumoral CD8^+^ T cells were similar in both groups. By contrast, tumor-specific tetramer-binding CD8^+^ T cell frequencies were increased ≥2-fold in all tissues apart from tumors (**Figure 2D-E**), suggestive of improved tumor-specific T cell development, survival, and/or proliferation. Notably, robust tetramer-binding T cell populations were recovered from *Tgfbr2* KO mice lacking macroscopic tumors, implying that *KPC*2a tumors were established and subsequently rejected following development of effective antitumor immunity. Similar to findings in the autochthonous *KPC* model, Klrg1^+^ T cell effector (T_EFF_) differentiation was increased by 2-fold in *Tgfbr2* KO hosts (**Figure 2F**). T cell exhaustion, assessed via co-expression of PD-1 and Lag3, was reduced in splenic and intratumoral tumor-specific T cells from *Tgfbr2* KO mice (**Figure 2G**). Stem-like progenitor (T_STEM_) cells, a T cell population that is believed to seed the effector and exhausted T cell compartments (46) were reduced by approximately 2-fold in all tissues of *Tgfbr2* KO mice (**Figure 2H**). However, increased frequencies of the *Tgfbr2* KO splenic and intratumoral T cells produced effector cytokines following restimulation with CB_101-109_ peptide (**Figure 2I**). Altogether, deletion of *Tgfbr2* in T cells augments tumor-specific CD8^+^ effector T cell generation and enhances their function, though these changes come at the expense of T_STEM_ formation and/or cell preservation/persistence.

**Figure 2.**
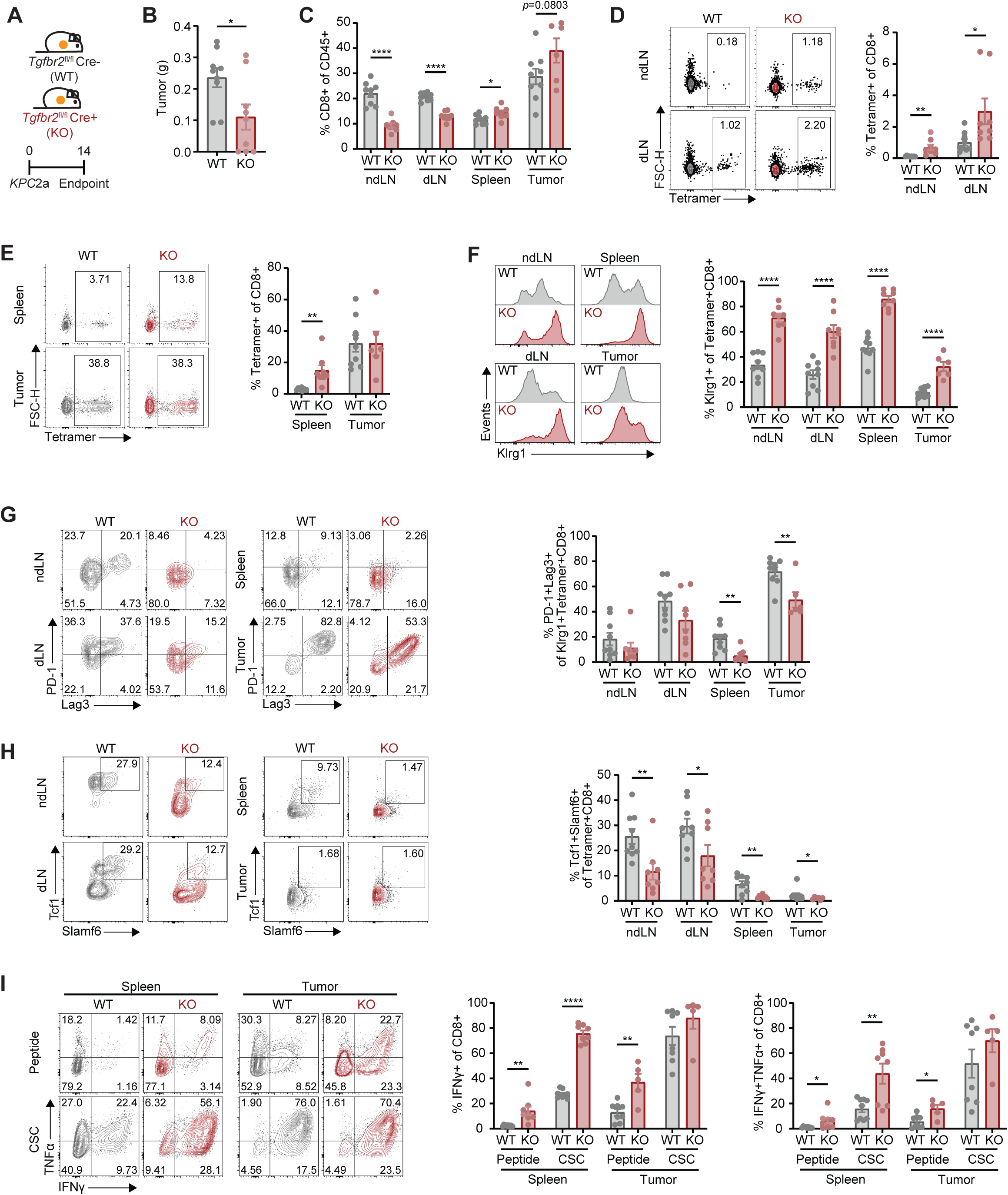
Abrogating T cell-intrinsic TGFβ improves endogenous neoantigen-specific T cells in orthotopic PDA. **A)** Schematic illustrating *KPC*2a (CBR^+^) PDA tumor implantation in Tgfbr2^Flox/Flox^ Cre^-^ (WT) or Tgfbr2^Flox/Flox^ Cre^+^ (KO) mice. **B)** Tumor weights on day 14. **C)** CD8^+^ T cell frequency of CD45^+^ cells in ndLN, dLN, spleen, and tumor. **D-E)** Representative plots (left) depicting CB_101-109_:H2D^b^ teramer^+^ frequency of CD8^+^ T cells in ndLN and dLN (D, right) or spleen and tumor (E, right). **F)** Representative histograms (left) of Klrg1^+^ frequency of tetramer^+^CD8^+^ T cells (right). **G)** Representative plots (left) of PD-1^+^Lag3^+^ frequency of tetramer^+^CD8^+^ T cells (right). **H)** Representative plots (left) of Tcf1^+^Slamf6^+^ frequency of tetramer^+^CD8^+^ T cells (right). **I)** Representative plots (left) of IFNγ^+^ (middle) or IFNγ^+^TNFα^+^ (right) frequency of tetramer^+^CD8^+^ T cells following restimulation with Msln_406-414_ or Cell Stimulation Cocktail (CSC). Results are compiled from two independent experiments. Dots are individual animals and n=8-9/group in all panels with mean ± SEM shown. Student’s t-test was used for comparisons in all panels.

### Vaccination improves therapeutic efficacy of 1045 TRex T cells in autochthonous and orthotopic PDA

PDA tumors generally exhibit low tumor mutational burden (TMB) and potent neoantigens may be rare (11–13). Thus, we returned to evaluate Msln as a clinically translational target in orthotopic models of PDA. We previously generated 1045 TCR exchange (TRex) mice, in which the 1045 TCR is integrated upstream of *Trac* while concomitantly deleting *Msln* to bypass tolerance (40, 43). As the donor 1045 TCR is located within the endogenous *Trac* locus, it is physiologically regulated (40, 43). To validate that donor T cells from TRex mice have antitumor activity in autochthonous PDA, we performed ACT with 5 x 10^6^ *in vitro* activated 1045 TRex T cells in *KPC* mice with established tumors. ACT was performed every two weeks with or without vaccination for a maximum of three doses (**Figure 3A**). The initial infusion was performed with agonistic anti-CD40 (αCD40), Msln_406-414_ peptide, and Poly(I:C) [TriVax (48)]. Subsequent infusions were administered with Msln_406-414_ peptide plus Poly(I:C) [BiVax (47)]. Previously employed cell-based vaccination was replaced with peptide-based vaccines given recent clinical success using similar approaches in PDA and other cancers (53–59). Median overall survival was prolonged in *KPC* mice receiving ACT with vaccination by nearly 5-fold compared to untreated mice (**Figure 3B**). Donor T cells were readily identified at endpoint (the time of euthanasia) by immunofluorescent microscopy from spleens of mice treated with ACT with or without vaccination. However, T cells were only identified in tumors of mice receiving ACT with vaccination (**Figure 3C**), suggesting that vaccination is essential for persistence of these donor T cells in treatment of PDA. To validate this observation, orthotopic tumors were established using our previously characterized *KPC*2 PDA line (41), and tumor-bearing mice were subsequently treated with 1045 TRex ACT with BiVax or TriVax (**Figure 3D**). Aligning with observations from autochthonous tumors in *KPC* mice, at day 14 donor T cells comprised just 1-10% of CD8^+^ T cells in all tissues of mice treated without vaccination, but were more than 40% of the CD8^+^ T cell compartment across tissues in mice receiving vaccination (**Figure 3E**). The inclusion of αCD40 in TriVax did not result in improved donor engraftment and/or expansion compared to BiVax. Thus, vaccination at the time of T cell transfer appears essential for ACT T cell persistence in both autochthonous and orthotopic models.

**Figure 3.**
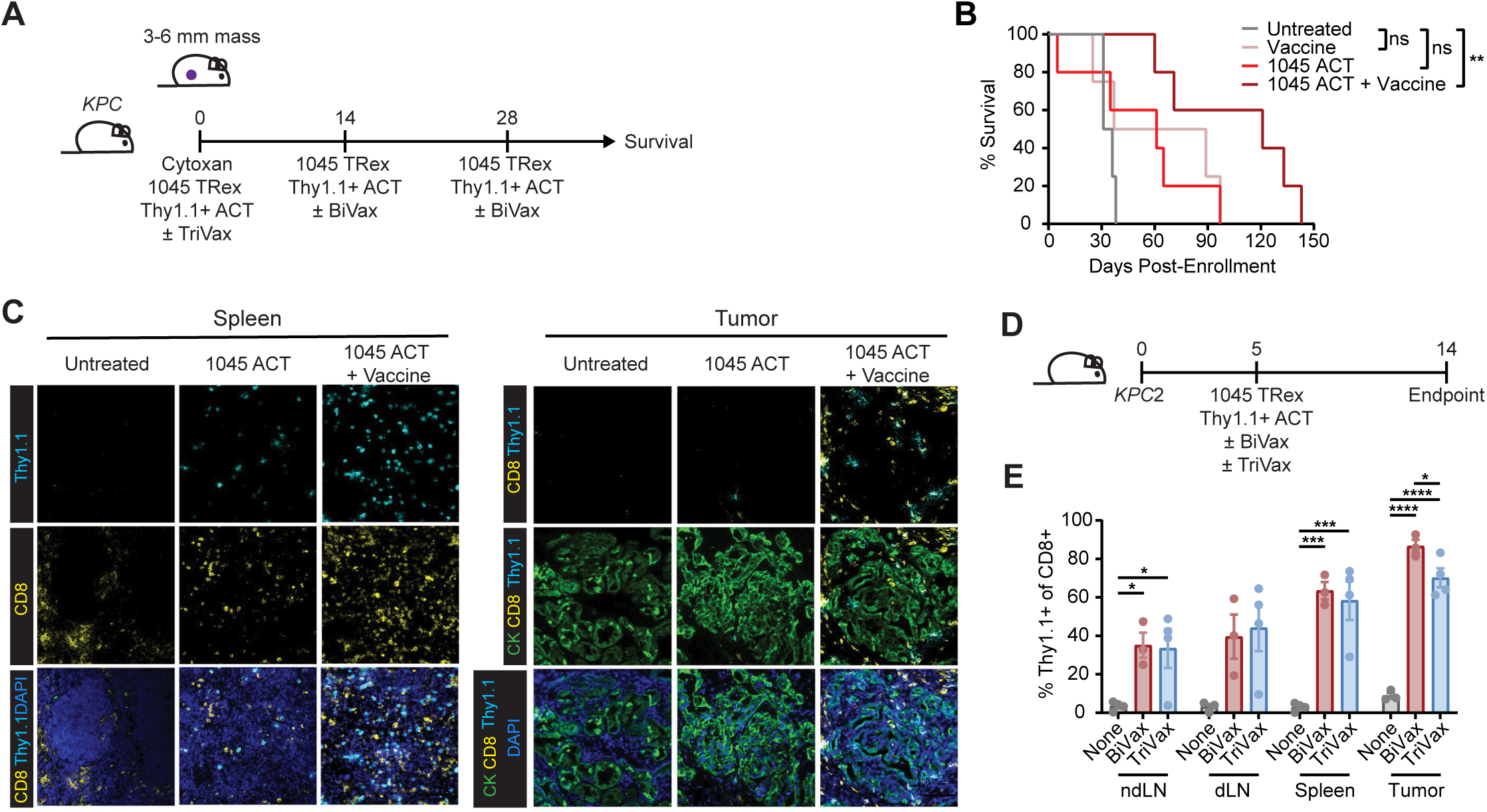
Vaccination improves therapeutic efficacy of 1045 TRex T cells in autochthonous and orthotopic PDA. **A)** Schematic illustrating biweekly ACT (5 × 10^6^ 1045 TRex WT Thy1.1^+^ T cells) with or without vaccination in tumor-bearing *KPC* mice pretreated with cyclophosphamide (120 mg/kg). **B)** Survival of *KPC* mice treated with ACT as depicted in (A). **C)** Representative images from immunofluorescent microscopy of spleens and tumors isolated from mice depicted in (A) at endpoint evaluating cytokeratin (CK), CD8, Thy1.1, and DAPI. **D)** Schematic of ACT (5 × 10^6^ 1045 TRex WT Thy1.1^+^ T cells) with or without vaccination in *KPC*2 tumor-bearing mice. **E)** Thy1.1^+^ donor frequency of CD8^+^ T cells from mice depicted in (D). Dots are individual animals and n=3-5/group in all panels with mean ± SEM shown. Cox regression was used for B and one-way ANOVA with Tukey posttest was used in E for comparisons.

### CRISPR/Cas9-mediated deletion of *Tgfbr2* in 1045 TRex T cells promotes effector T cell accumulation and interactions with cDC1s in orthotopic KPC tumors

We next queried if 1045 TRex T cell antitumor responses were augmented in orthotopic PDA by targeted deletion of *Tgfbr2* using a CRISPR/Cas9 based approach (generating “1045 TRex *Tgfbr2*^CRISPR^” T cells). A mixture of two guide RNAs (gRNA) were used for targeted deletion of *Tgfbr2* in 1045 TRex cells and resulted in >50% editing efficiency (**Figure 4A**). TGFβR2 loss was verified using flow cytometry (**Figure 4B**). As deletion of *Tgfbr2* improved effector differentiation of donor T cells (Figure 1F-H), we hypothesized that TGFβ signaling limits functional interactions between 1045 TRex T cells and type I dendritic cells (cDC1s), given these specialized antigen-presenting cells (APCs) are required for differentiation and sustained antitumor function following ACT (44). Indeed, cDC1s are uniquely efficient at cross-presenting cell-associated tumor antigen (44, 60) and therefore essential for CD8^+^ T cell function. To interrogate this hypothesis, we orthotopically implanted mScarlet^+^ *KPC*2 cells into Xcr1-Venus mice, which permitted visualization of tumor cells (red) and cDC1s (yellow). Seven days after tumor establishment, we performed ACT with 1045-eGFP WT or *Tgfbr2*^CRISPR^ T cells using our protocol that included transient lymphodepletion with cyclophosphamide and vaccination with TriVax (**Figure 4C**). Seven days after transfer, we performed 2-photon and second harmonic imaging (SHG) of donor T cells and cDC1s (**Figure 4D**). Intratumoral 1045-eGFP *Tgfbr2*^CRISPR^ T cells exhibited decreased motility with reduced mean distance traveled and displacement (**Figure 4E-F**). However, mean directional change was increased and confinement ratio was decreased. Thus, cell-cell interactions in 1045-eGFP Tgfbr2^CRISPR^ T cells appeared to be prolonged. cDC1 motility was increased in mice that received 1045-eGFP *Tgfbr2*^CRISPR^ T cells, as exhibited by increased distance traveled, displacement, and speed (**Figure 4E, G**). Together, the data support a model in which loss of *Tgfbr2* in T cells may increase cDC1 environmental sampling, but interactions between donor T cells and cDC1s, when they do occur, are prolonged. These dynamics might explain increased differentiation of donor T cells toward an effector state.

**Figure 4.**
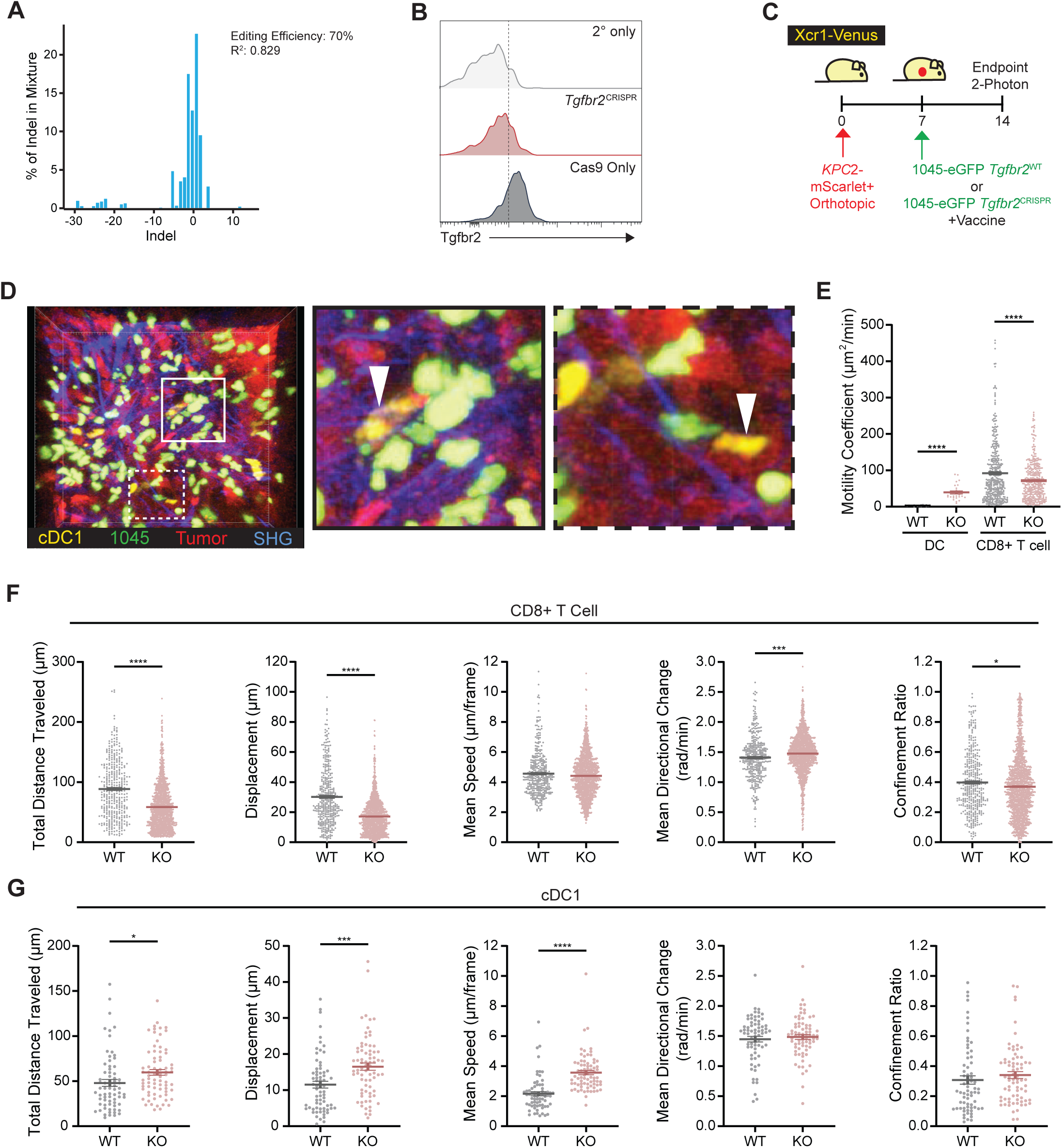
CRISPR/Cas9-mediated deletion of *Tgfbr2* in 1045 TRex T cells promotes effector T cell accumulation and interactions with cDC1s in orthotopic KPC tumors. **A)** ICE analysis from representative CRISPR/Cas9-mediated deletion of *Tgfbr2* in 1045 TRex T cells. **B)** Histograms from representative preparation of 1045 TRex *Tgfbr2*^CRISPR^ cells with secondary antibody only (upper histogram) and Cas9 only edited cells (lower histogram) as controls. **C)** Schematic showing ACT with 5 × 10^6^ 1045-eGFP *Tgfbr2*^WT^ (WT) or *Tgfbr2*^CRISPR^ T cells in Xcr1-Venus mice harboring *KPC*2 mScarlet^+^ tumors with TriVax for two-photon evaluation. **D)** Representative still frame from two-photon video derived from an animal treated with 1045-eGFP *Tgfbr2*^CRISPR^ ACT. Insets (white squares) at higher magnification depict cDC1 cells (white arrowheads) in close proximity to 1045-eGFP *Tgfbr2*^CRISPR^ T cells. **E)** Motility coefficients from cDC1 and 1045-eGFP *Tgfbr2*^WT^ or *Tgfbr2*^CRISPR^ T cells. **F-G)** Total distance traveled, displacement, mean speed, mean directional change, and confinement ratio of T cells (F) or cDC1s (G) in mice treated with 1045-eGFP *Tgfbr2*^WT^ or *Tgfbr2*^CRISPR^ (“KO”) ACT. Two-photon measurements are derived from n=3/group with mean ± SEM shown. Student’s t-test was used for comparisons in all panels.

### Genetic deletion of *Tgfbr2* in 1045 TRex T cells promotes enhanced antitumor T cell function *in vitro*

The above CRISPR/Cas9 approach was limiting due to incomplete *Tgfbr2* editing efficiency, making long-term assessment after ACT difficult to interpret. Therefore, to more robustly assess the impact of abrogating TGFβ signaling in T cell antitumor efficacy, we crossed 1045 TRex mice to the *Tgfbr2*^Flox/Flox^ x d*Lck*-Cre strain employed in Figure 1. CD8^+^ T cell frequencies in the resultant mice were identical in 1045 TRex *Tgfbr2*^Flox/Flox^ Cre^-^ (TRex WT) and Cre^+^ (TRex KO) mice, though CD4^+^ T cells were more prevalent in TRex KO mice (**Supplemental Figure 1A-C**). 1045 TCR frequency, assessed by measuring Vβ9, was reduced by approximately 25% in TRex KO compared to WT mice (**Supplemental Figure 1D**). >80% of Vβ9^+^CD8^+^ T cells present with spleen and LN of both TRex WT and KO mice were naïve (CD44^-^CD62L^+^) (**Supplemental Figure 1E-F**). In contrast, Vβ9^-^CD8^+^ T cells, which include polyclonal and potentially self-reactive T cells, formed central memory (CD44^+^CD62L^+^, T_CM_) and T_EFF_ (CD44^+^CD62L^-^) phenotypes more readily. Klrg1 and Cx3cr1, further markers associated with T_EFF_ differentiation, were only readily detected on splenic Vβ9^-^CD8^+^ T cells of both strains and were increased 2-fold in TRex KO mice (**Supplemental Figure 1G-I**). Thus, loss of TGFβ signaling in 1045 TRex KO mice does not result in enhanced 1045 T cell recognition of the low endogenous levels of Msln in healthy tissues.

The functions of 1045 TRex WT and KO cells were then measured using *in vitro* assays. Naïve TRex KO cells exhibited enhanced proliferation following *in vitro* activation with titrating amounts Msln_406-414_ peptide (**Figure 5A-D**). Production of effector cytokines and GzmB were quantified in restimulated TRex T cells (**Figure 5E**).

**Figure 5.**
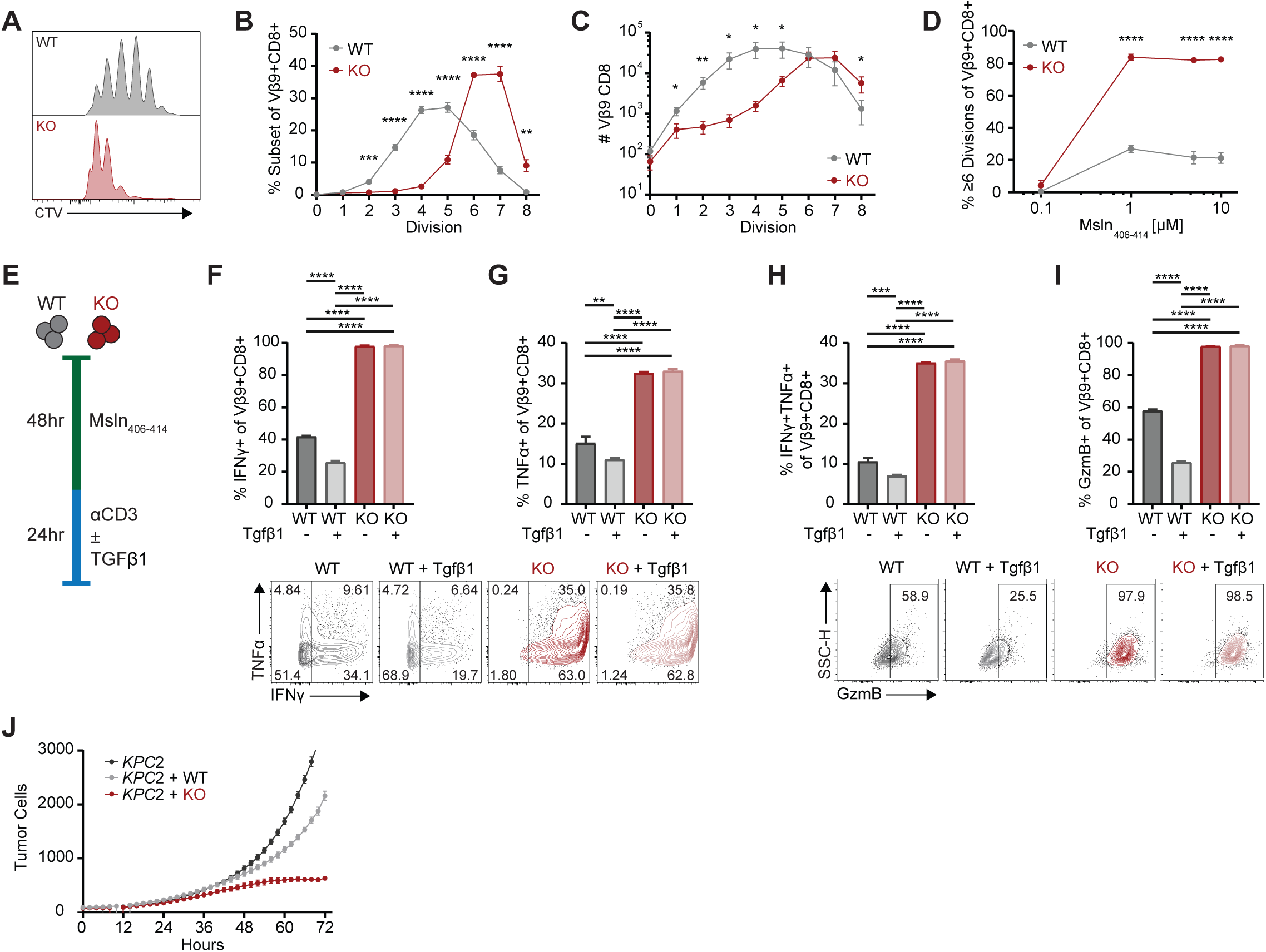
Genetic deletion of *Tgfbr2* in 1045 TRex T cells promotes enhanced antitumor T cell function *in vitro*. **A)** Representative histograms of proliferation of 1045 TRex WT or KO T cells stimulated with 1 μg/mL Msln_406-414_ for 72 hours as assessed by CellTrace Violet (CTV) dilution. **B)** Frequency of 1045 TRex WT or KO T cells stimulated with 1 μg/mL Msln_406-414_ by number of divisions. **C)** Number of 1045 WT or KO T cells stimulated with 1 μg/mL Msln_406-414_ by number of divisions. **D)** Frequency of 1045 TRex WT or KO T cells stimulated with 0.1-10 μg/mL Msln_406-414_ reaching ≥6 divisions. **E)** Schematic illustrating priming and restimulation with or without 20 ng/mL Tgfβ1 of 1045 TRex WT or KO T cells to assess for cytokine production. **F-I)** IFNγ^+^ (F), TNFα^+^ (G), IFNγ^+^TNFα^+^ (H), or GzmB^+^ (I) frequency of 1045 TRex WT or KO T cells following restimulation as in (E). Representative histograms are shown in lower panels. **J)** *KPC*2 NIR^+^ tumor cell number following co-culture with 1045 TRex WT or KO T cells at a 5:1 E:T. All panels show mean ± SD from experimental triplicates and are representative of at least three independent experiments. Student’s t-test was used in B-D and One-way ANOVA with Tukey posttest was used in F-I for comparisons.

IFNγ, TNFα, and GzmB-producing 1045 TRex KO cells were increased 2 to 3-fold over 1045 TRex WT cells (**Figure 5F-I**), aligning with initial studies on TGFβ in T cells (38). The addition of Tgfβ1 during restimulation significantly blunted cytokine production in TRex WT cells by 2-fold. Finally, cytotoxicity was assessed and revealed increased lysis of *KPC*2 cells co-cultured with 1045 TRex KO effectors, though some *KPC*2 target cells were able to persist (**Figure 5J**). These results reaffirm that TGFβ limits proliferation, cytokine production, and cytotoxicity in CD8^+^ T cells, likely in part by attenuating TCR signaling (33, 34).

### Efficacy of adoptively transferred TCR-engineered T cells is limited in orthotopic PDA

To determine if abrogating TGFβ signaling could increase the efficacy of 1045 TRex ACT, *KPC*2 orthotopic-bearing mice were treated with TRex WT or KO ACT 5 days after orthotopic *KPC*2 implantation (**Figure 6A**). Recipient mice received lymphodepletion followed by ACT with BiVax and were assessed 15 days after tumor implantation (10 days after ACT). TRex WT and KO ACT similarly reduced *KPC*2 tumor weights by a modest ∼30% on day 15 compared to tumors from control mice (**Figure 6B**). Akin to observations in autochthonous PDA, frequencies of 1045 TRex KO donor CD8^+^ T cells were increased 2-fold compared to WT counterparts in ndLNs, dLNs, spleens, and tumors of treated mice (**Figure 6C-D**). Klrg1^+^ T_EFF_ differentiation was increased in TRex KO compared to WT cells (**Figure 6E-F**). However, this differentiation phenotype occurred in <10% of KO donor cells – markedly reduced as compared to evaluations in prior models (**Figures 1F-H and 2F**).

**Figure 6.**
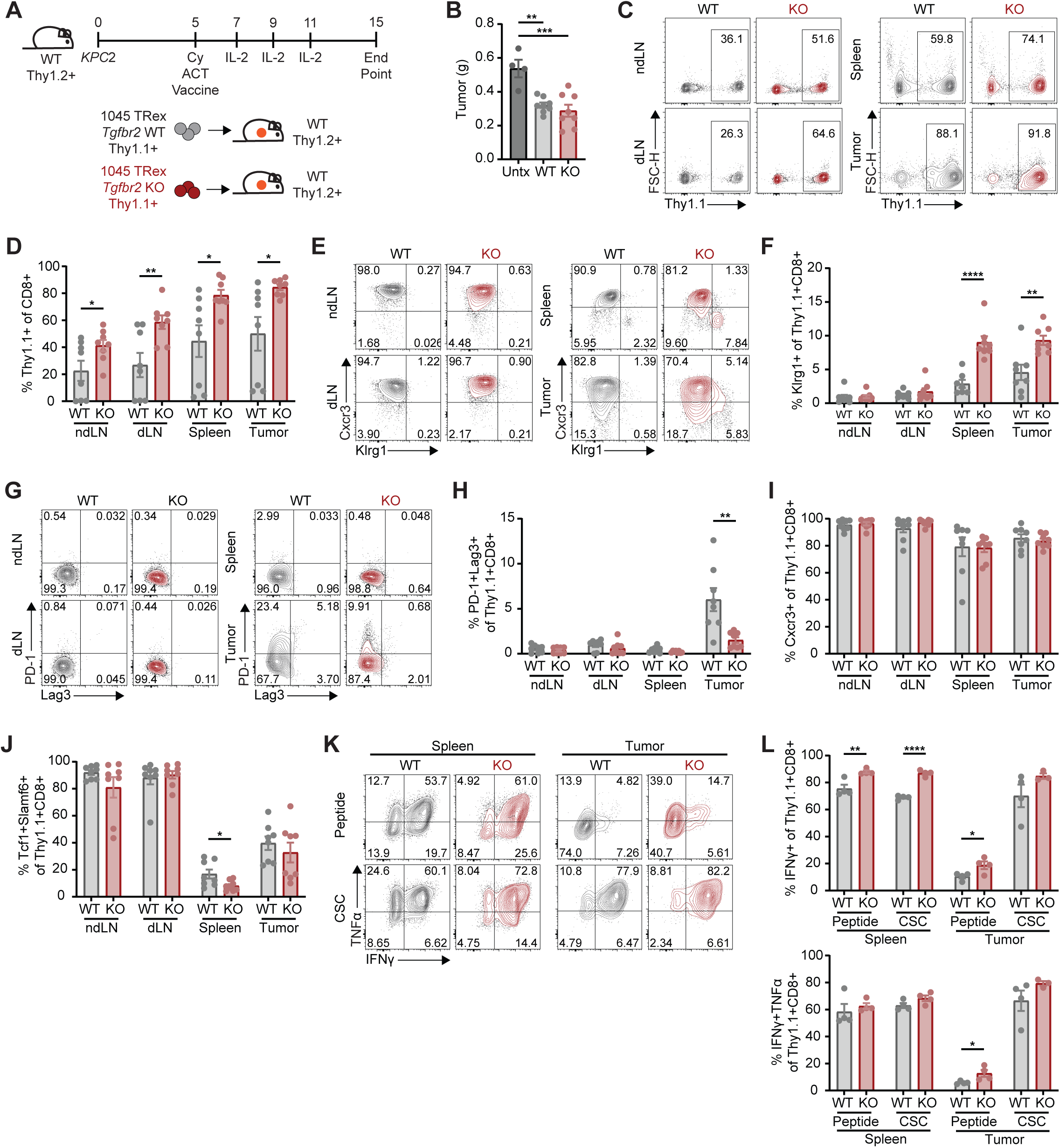
Efficacy of adoptively transferred TCR-engineered T cells is limited in orthotopic PDA. **A)** Schematic showing 1045 TRex WT or KO ACT (5 × 10^6^ T cells) plus BiVax in *KPC*2 tumor-bearing hosts pretreated with cyclophosphamide (120 mg/kg). **B)** Tumor weights on day 15. **C-D)** Representative plots (C) of Thy1.1^+^ 1045 TRex WT or KO donor frequency of CD8^+^ T cells in ndLN, dLN, spleen, and tumor (D). **E)** Representative plots of Cxcr3 and Klrg1 on 1045 TRex WT or KO donor cells. **F)** Klrg1^+^ frequency of 1045 TRex WT or KO donor cells. **G-H)** Representative plots (G) depicting PD-1^+^Lag3^+^ frequency of 1045 TRex WT or KO donor cells (H). **I)** Cxcr3^+^ frequency of 1045 TRex WT or KO donor cells [see representative plots in (E)]. **J)** Tcf1^+^Slamf6^+^ frequency of 1045 TRex WT or KO donor cells. **K-L)** Representative plots (K) depicting IFNγ^+^ (L, upper) or IFNγ^+^TNFα^+^ (L, lower) frequency of 1045 TRex WT or KO donor cells following restimulation with Msln_406-414_ or CSC. Results are compiled from two independent experiments. Dots are individual animals and n=4-8/group in all panels with mean ± SEM shown. One-way ANOVA was used in B with Student’s t-test used for comparisons in all remaining panels.

Additionally, T cell exhaustion quantified by co-expression of PD-1 and Lag3 was also limited and detected in <10% of donor 1045 TRex WT and KO T cells (**Figure 6G-H**). Cxcr3, a chemokine receptor enriched in memory T cell populations (45), was expressed by nearly all TRex WT and KO donor T cells (**Figure 6E, I**). T_STEM_ readily developed within LNs and comprised ∼90% of both groups of donor T cells (**Figure 6J**). Further, approximately 40% of intratumoral donor WT and KO T cells formed T_STEM_, substantially increased compared to intratumoral levels observed in *Tgfbr2*^Flox/Flox^ x d*Lck*-Cre mice challenged with *KPC*2a (Figure 2H). Modestly increased IFNγ production was observed in splenic TRex KO cells following restimulation with Msln_406-414_ or a non-specific stimulus (**Figure 6K-L**). A similar modest increase in IFNγ production was observed in restimulated intratumoral TRex KO donor cells. However, ≥80% of both TRex WT and KO intratumoral donor cells exhibited poor reactivity to cognate antigen. Collectively, these data indicate a fundamental defect in 1045 TRex T cells when encountering *KPC*2 tumor cells in an orthotopic setting, which ultimately resulted in a quiescent T state with poor effector function that was not overcome by loss of TGFβ signaling.

TGFβ has established roles in T cell memory formation, particularly tissue-resident memory (T_RM_) (61, 62). Curiously, memory phenotypes formed similarly in *KPC*2-bearing mice treated with either TRex WT or KO ACT. To more directly examine how homeostatic levels of TGFβ influence memory formation following ACT, we evaluated T cells at memory timepoints in non-tumor-bearing mice. Thy1.1^+^/1.1^+^ TRex WT and Thy1.1^+^/1.2^+^ TRex KO T cells were co-transferred at a 1:1 ratio in healthy mice using our ACT protocol with cyclophosphamide-based lymphodepletion and BiVax (**Supplemental Figure 2A-B**). Donor cells were tracked longitudinally in the blood for 6 weeks. Circulating TRex KO T cells were enriched 10-to 20-fold and comprised around 80% of CD8^+^ T cells within hosts during the 6 weeks immediately following transfer (**Supplemental Figure 2C-F**). CD127 and CD62L were used to delineate T_CM_ (CD127^+^CD62L^+^), effector memory (T_EM_, CD127^+^CD62L^-^), and T_EFF_ (CD127^-^CD62L^-^) (**Supplemental Figure 2G-J**). TRex KO T cells formed reduced frequencies of T_CM_ and increased T_EM_ and T_EFF_ compared to TRex WT T cells, though these differences were modest but statistically significant. Thus, 1045 TRex cells form T_CM_ in the absence of TGFβ.

T_RM_ were enumerated within a broad array of tissues [inguinal LN, spleen, pancreas, lung, liver, and intraepithelial small intestine (SI-IEL)] 8 weeks after co-transfer using intravenous labeling of CD45^+^ leukocytes. Similar to circulating T cell findings, extravascular TRex KO T cells were approximately 10-fold more prevalent in all tissues apart from SI-IEL (**Supplemental Figure 3A-B**). T_RM_-associated markers, including CD69, CD103, CD49a, and Cxcr6, were interrogated on donor cells within each tissue compartment, alongside endogenous cells within SI-IEL, which served as a positive control given nearly all CD8^+^ T cells within SI-IEL are T_RM_ (63). CD103 was undetectable on extravascular TRex KO T cells (**Supplemental Figure 3C**), which was predictable given TGFβ directly and indirectly regulates CD103 (61, 62). CD69 expression on extravascular TRex WT and KO T cells was nearly identical in all tissues apart from SI-IEL, where a nearly 2-fold reduction in TRex KO T cells was present (**Supplemental Figure 3D**). Similar trends were observed in CD49a and Cxcr6 (**Supplemental Figure 3E-F**). CD69^+^CD49a^+^ and CD49a+Cxcr6+ co-expression was similar in both TRex WT and KO T cells in most tissues, again except for SI-IEL (**Supplemental Figure 3G-H**).

CD69^+^Cxcr6^+^ and co-expression of all three markers were reduced in TRex KO T cells by ∼30-50% (**Supplemental Figure 3I-J**). Frequencies of all T_RM_ markers were similarly reduced on TRex KO T cells in SI-IEL. Collectively, these findings indicate that memory T cell differentiation readily occurs in the absence of TGFβ, and T_CM_ and T_RM_ formation are only modestly impaired in the experimental setting of ACT.

### Msln processing restricts TCR signaling while TGFβ-dependent and independent mechanisms suppress IFNγ production *in vitro*

Due to the lack of effector and exhausted T cell differentiation in orthotopic *KPC*2 tumors (Figure 6E-H), we hypothesized that Msln epitope processing and/or presentation limits 1045 T cell recognition of *KPC* tumor cells. We first confirmed that three *KPC* PDA lines (*KPC*2, *KPC*271, *KPC*451) derived from independent *KPC* mice express *Msln* transcript (**Supplemental Figure 4A**). The levels of *Msln* transcripts varied somewhat but were detected in all *KPC* lines. Intracellular and membrane-bound Msln was also measured (**Supplemental Figure 4B**). Though membrane-bound Msln was variable, perhaps partially owing to variable Msln cleavage and shedding (64), intracellular Msln was similar in all three *KPC* cell lines. H2D^b^ was expressed and upregulated following IFNγ treatment in all *KPC* cell lines (**Supplemental Figure 4C**). *Tgfβ1* transcripts were assessed and found to be robustly expressed (**Supplemental Figure 4D**). Latency activated peptide (LAP), a prodomain which complexes with TGFβ and is cleaved to generate active TGFβ (65), was used as a surrogate to evaluate TGFβ in *KPC*2 tumors (**Supplemental Figure 4E-F**). LAP was robustly identified within *KPC*2 tumor cell rich zones, suggesting that Tgfβ1 is primarily derived from PDA tumor cells *in vivo*. Thus, Msln, MHC-I, and Tgfβ1 are expressed and/or produced by diverse *KPC* cell lines.

To interrogate Msln antigen processing and presentation, *KPC*2 was engineered to produce Msln_406-414_ peptide (*KPC*2^Msln-ER^) using a previously established strategy (**Supplemental Figure 4G**) (42). The approach targets the antigen peptide to the endoplasmic reticulum (ER), where it can be loaded on MHC-I, and therefore allows for bypass of antigen processing (42). *KPC*2^Msln-ER^, *KPC*2, *KPC*271, and *KPC*451 were co-cultured with primed 1045 TRex WT or KO T cells for 24 hours followed by measurement of activation markers associated with TCR signaling (CD25, CD69) along with effector molecule (GzmB, IFNγ) production (**Figure 7A**). Target cells were pretreated with IFNγ (to upregulate MHC-I) and/or soluble Msln_406-414_ peptide as positive controls. CD25^+^CD69^+^co-expression occurred in only 10-20% of 1045 TRex WT T cells co-cultured with the various *KPC* cell lines (**Figure 7B**). In contrast, ∼40% of 1045 TRex KO T cells were CD25^+^CD69^+^, suggesting improved tumor cell antigen recognition and sensitivity of TCR signaling in the absence of *Tgfbr2*. Overexpression of Msln_406-414_ in *KPC*2^Msln-ER^ cells resulted in modest but significant increases in tumor antigen recognition when compared to control *KPC*2 cells. Pretreating *KPC* tumor cells with IFNγ to increase cell-surface peptide:MHC-I approximately doubled CD25^+^CD69^+^ frequency in 1045 TRex WT T cells, though only modest increases were observed in 1045 TRex KO T cells. Together, these findings suggest that loss of TGFβ augments TCR signaling in response to limiting amounts of antigen. Controls consisting of peptide-pulsed *KPC* cells with or without IFNγ resulted in CD25^+^CD69^+^ frequencies of 50-80% in both 1045 TRex WT and KO T cells. Thus, both Msln antigen processing and MHC-I limit T cell recognition of *KPC* tumor cells. Similar trends were observed in co-cultures when examining GzmB (**Figure 7C**). However, <10% of 1045 TRex WT and KO T cells produced IFNγ, even when target cells were both pulsed with cognate antigen and pretreated to induce MHC-I. Therefore, tumor cell-intrinsic mechanisms that span diverse *KPC* tumor lines suppress IFNγ, even when T cell antigen recognition is optimized.

**Figure 7.**
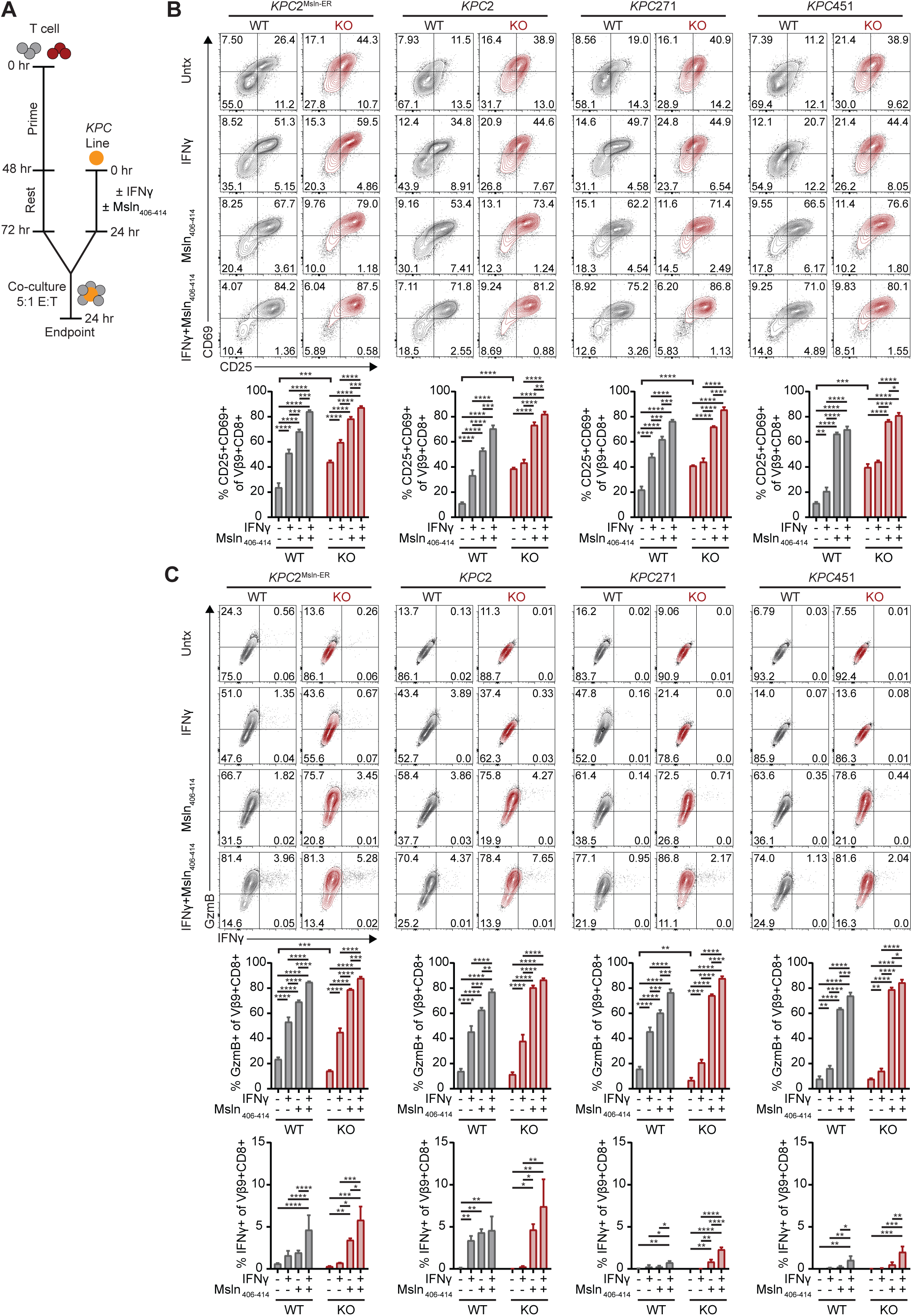
Msln processing restricts TCR signaling while TGFβ-dependent and independent mechanisms suppress IFNγ production *in vitro*. **A)** Schematic showing preparation of 1045 TRex WT and KO T cells along with *KPC* tumor cell lines for co-culture. Briefly, T cells were primed with Msln_406-414_ for 48 hours followed by a 24 hour rest period. Tumor cells were plated with or without IFNγ (100 μg/mL) for 24 hours and/or pulsed with Msln_406-414_ for one hour prior to co-culture. Prepared T cells were plated at a 5:1 E:T and co-cultured for 24 hours. **B)** Representative plots (above) from co-cultures prepared with *KPC*2^Msln-ER^, *KPC*2, *KPC*271, and *KPC*451 depicting CD25^+^CD69^+^ frequency of 1045 TRex WT and KO T cells (lower). **C)** Representative plots (upper) depicting GzmB^+^ (middle) or IFNγ^+^ (lower) frequency of 1045 TRex WT and KO T cells. All panels show mean ± SD from experimental triplicates and are representative of at least three independent experiments. One-way ANOVA with Tukey posttest was used for comparisons in all panels.

### Enforcing Msln antigen presentation restores *in vivo* antitumor function in PDA

As suboptimal Msln antigen processing partially restricted T cell recognition of *KPC* tumor cells (Figure 7), we then examined efficacy of 1045 TRex ACT in mice with orthotopic *KPC*2^Msln-ER^ tumors. Mice with established tumors were initially treated with cyclophosphamide followed by ACT without vaccination 6 days after orthotopic *KPC*2^Msln-ER^ tumor implantation (**Figure 8A**). Tumor growth, assessed by ultrasound, was reduced in mice treated with ACT and further suppressed when 1045 TRex KO cells were employed (**Figure 8B-C**).

**Figure 8.**
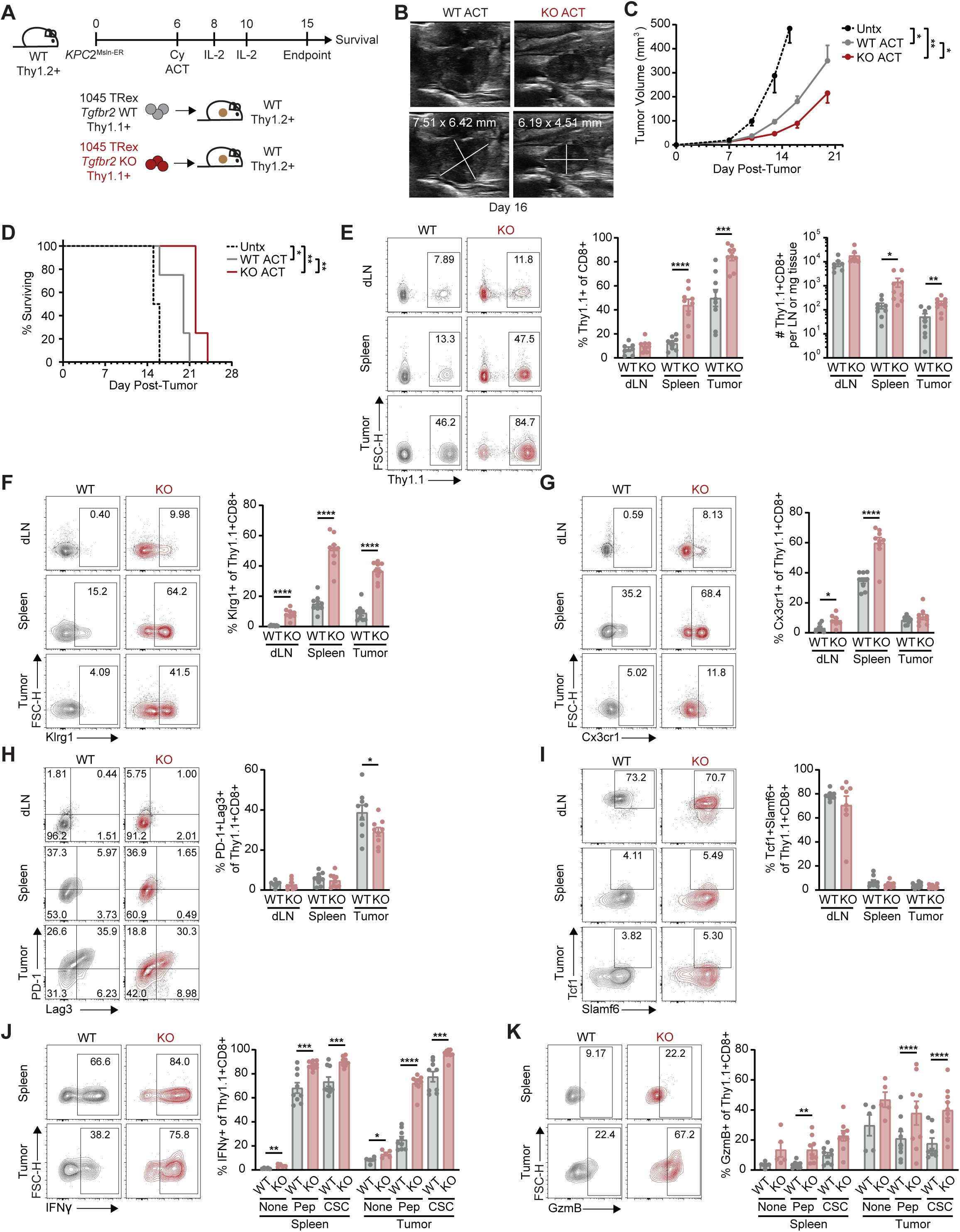
Enforcing Msln antigen restores *in vivo* antitumor function in PDA. **A)** Schematic illustrating 1045 TRex WT or KO ACT (5 × 10^6^ T cells) in *KPC*2^Msln-ER^ tumor-bearing hosts pretreated with cyclophosphamide (120 mg/kg). **B)** Representative ultrasound images from mice treated with 1045 TRex WT or KO ACT on day 16. **C)** Tumor volumes from control or treated mice assessed using ultrasound. **D)** Survival of control and treated mice. **E)** Representative plots (left) depicting Thy1.1^+^ donor frequency of CD8^+^ T cells (middle) or Thy1.1^+^ donor cell number per LN or mg of tissue (right). **F)** Representative plots (left) depicting Klrg1^+^ frequency of 1045 TRex WT or KO donor cells (right). **G)** Representative plots (left) depicting Cx3cr1^+^ frequency of 1045 TRex WT or KO donor cells (right). **H)** Representative plots (left) depicting PD-1^+^Lag3^+^ frequency of 1045 TRex WT or KO donor cells (right). **I)** Representative plots (left) depicting Tcf1^+^Slamf6^+^ frequency of 1045 TRex WT or KO donor cells (right). **J)** Representative plots (left) depicting IFNγ^+^ frequency of unstimulated 1045 TRex WT or KO donor cells or in cells restimulated with Msln_406-414_ or CSC (right). **K)** Representative plots (left) depicting GzmB^+^ frequency of unstimulated 1045 TRex WT or KO donor cells or in cells restimulated with Msln_406-414_ or CSC (right). C-D are representative of two independent experiments and contain n=4/group. E-K are compiled from two independent experiments with dots representing individual animals, n=8/group, and mean ± SEM shown. One-way ANOVA with Tukey posttest was used in C, Cox regression was used in D, and Student’s t-test was used in all remaining panels for comparisons.

Survival was also significantly prolonged by 1045 TRex KO ACT compared to WT ACT, though rapid tumor growth was observed in all treated mice after 14 days (**Figure 8D**), suggesting either development of resistance or loss of T cell-mediated antitumor efficacy.

T cell persistence and phenotypes were further quantified on day 15 (9 days after transfer). By frequency, 1045 TRex KO T cells were enriched by ∼2-to 3-fold compared to 1045 TRex WT T cells in the spleens and tumors of treated mice (**Figure 8E**). By numbers, approximately 10-fold and 5-fold more TRex KO T cells were recovered from spleens and tumors of mice treated with 1045 TRex KO ACT, respectively. Aligning with our prior findings, abrogating TGFβ signaling resulted in >4-fold increases in Klrg1^+^ T_EFF_ differentiation (**Figure 8F**). An additional effector memory marker, Cx3cr1, was also increased in 1045 TRex KO T cells present within dLN and spleens (**Figure 8G**). In contrast to mice with *KPC*2 orthotopic tumors treated with 1045 TRex ACT, ∼40% of 1045 TRex WT T cells within *KPC*2^Msln-ER^ tumors developed evidence of exhaustion (PD-1^+^Lag3^+^) (**Figure 8H**). Intratumoral 1045 TRex KO T cells exhibited slightly reduced (∼30%) co-expression of exhaustion markers. T_STEM_ formation was similar in both 1045 TRex WT and KO T cells, and splenic and intratumoral T_STEM_ accounted for <10% of total donor cells (**Figure 8I**). Expectedly, unstimulated splenic donor T cells produced minimal IFNγ, though the frequency of cytokine-producing cells was slightly increased in TRex KO T cells (**Figure 8J**). 70% of splenic 1045 TRex WT T cells retained functionality when restimulated with Msln_406-414_ peptide or potent non-specific stimuli. This frequency increased to ∼90% in 1045 TRex KO cells shielded from TGFβ. In contrast, intratumoral 1045 TRex WT T cells exhibited signs of dysfunction, and only one-third of cells produced IFNγ in response to Msln_406-414_ peptide. However, 80% of intratumoral 1045 TRex KO cells retained the ability to produce IFNγ. Though variable, similar trends were observed when evaluating GzmB production, which increased nearly 2-fold in intratumoral 1045 TRex KO cells restimulated with cognate peptide (**Figure 8K**). Altogether, these findings reveal that protecting tumor-specific T cells from TGFβ confers improved effector differentiation and function, and so long as sufficient tumor epitope is presented, may bypass the T cell persistence advantage provided by vaccination.

## Discussion

Here, we establish the impact of abrogating TGFβ signaling in TCR-engineered CD8^+^ T cells specific for the tumor-associated antigen, Msln. T cells with deletion of *Tgfbr2* exhibited increased proliferation, cytokine production, and cytotoxic capacity *in vitro*. These attributes combined with enriched effector differentiation were consistently observed across several PDA *in vivo* tumor models. Specifically, in heterogeneous autochthonous tumors, Msln-specific T cells unresponsive to TGFβ outcompeted T cells with intact TGFβ signaling and more efficiently adopted effector phenotypes. Similarly, endogenous development of neoantigen-specific T cells was augmented when TGFβ signaling was impaired and formed effector phenotypes with greater frequency. The enhanced fitness linked to protection from TGFβ ultimately slowed PDA progression and prolonged survival in orthotopic models so long as sufficient Msln epitope was presented. These findings substantiate that TGFβ is a prominent immunosuppressive factor in PDA that limits the development of effective antitumor immunity, particularly in the context of ACT.

While diverse immunosuppressive mechanisms such as Foxp3^+^ regulatory T cells (T_REG_) and cancer-associated fibroblasts (CAFs) are enriched in tumors, many of the suppressive mechanisms converge on TGFβ. As such, protecting T cells from suppressive TGFβ has been pursued using several approaches. Dominant-negative (DN) forms of TGFβR2, initially developed in 1993 (66, 67), have been evaluated extensively in hematologic, gastrointestinal, genitourinary, brain, and other malignancies using TCR-engineered T cell ACT, CAR-T, and other therapies (68–80). Encouraging findings have led to early phase clinical trials in Hodgkin lymphoma and prostate cancer (81, 82). Notably, DN TGFβR2 T cells can drive lymphoproliferative disorders (83), raising long-term safety concerns when using this engineering strategy in cellular therapies. Thus, genetic deletion of *Tgfbr2* in T cells, which does not appear to be associated with oncogenesis, has also been explored in several cancer models (84–86). More recent novel approaches to target detrimental TGFβ signaling in T cells include switch receptors (87, 88) and improved DN constructs (89). Our findings align with and add to this collective knowledge, and support that protecting tumor-specific T cells from TGFβ augments their function even in PDA. Enhanced effector differentiation in TGFβ insensitive T cells may be related to prolonged interactions with cDC1s within tumors, which we observed in orthotopic PDA. Such findings correspond with prior studies which have demonstrated that TGFβ mitigates intratumoral T cell infiltration, resulting in exclusion of T cells and accumulation within the periphery of TGFβ-dense tumors (90, 91). Our approach herein also appears to be safe and tolerable, as no evidence of toxicity was observed in any of the treated mice reported above.

Our study supports that memory T cells readily develop in the absence of TGFβ when using an ACT protocol. Indeed, findings in DN Tgfbr2 and Tgfbr2 KO studies have generated conflicting results, with some reporting increased memory T cell formation (74, 79, 80, 84), while others have reported loss of or no change in memory potential (76, 77, 83). TGFβ has been particularly well studied in the generation of T_RM_ (61, 62). While TGFβ clearly contributes to T_RM_ formation, recent elegant studies have shown that TGFβ is only required for T_RM_ in certain tissues and contexts (92). Specifically, while TGFβ is absolutely required for cutaneous T_RM_ (the focus of most T_RM_ evaluations), it appears to also contribute to T_RM_ within SI, but may be completely dispensable in other tissues such as liver (92). We found that circulating central memory cells readily form in T cells insensitive to TGFβ, at least in the context of ACT. Additionally, T_RM_ were maintained or only modestly reduced in LNs, spleen, pancreas, lung, and liver, suggesting that TGFβ has only a minor role in establishing T_RM_ within these tissues. This apparent tissue residency occurred in a CD103-independent manner, given *Tgfbr2*^-/-^ T cells completely lacked CD103. As these memory experiments were performed in healthy mice, they do not incorporate any systemic immunosuppressive effects that tumors including PDA may promote. However, as *Tgfbr2*^-/-^ 1045 TRex T cells readily accumulated in tumors despite a lack of CD103 upregulation, the data also support that engineered T cell accumulation and effector function in PDA is CD103 independent. Nevertheless, future studies using less aggressive models of PDA and perhaps curative therapies are required to fully understand T_RM_ formation in ACT for PDA and other malignancies.

Donor T cell expansion and persistence is a major limitation in ACT which hinders efficacy (93–98). We demonstrate that vaccination was essential to promote ACT cell product engraftment and expansion in both autochthonous and orthotopic PDA models. These findings suggest that delivery of antigen in the context of inflammatory cues provided by adjuvant instructs tumor-specific T cells to engraft and expand during ACT. The benefits of vaccination further suggests that Msln antigen presentation is limiting in PDA. This hypothesis was further validated by studying Msln-targeted ACT in orthotopic models of PDA where efficacy of ACT using T cells subject to TGFβ signaling and those protected from it both resulted in poor efficacy. Phenotypic analysis of donor T cells revealed limited effector or exhausted differentiation with associated memory phenotypes that were not rescued by protecting T cells from suppressive TGFβ. Though dysfunction, as assessed by cytokine production upon restimulation, was mostly restored through deletion of *Tgfbr2*, the lack of efficacy of ACT with *Tgfbr2*^-/-^ cells strongly suggested loss or impairment in Msln processing and/or presentation by pancreatic tumor cells. By assessing multiple *KPC* cell lines, we determined that while Msln was abundantly expressed in tumor epithelial cells, T cells poorly recognized the target cells. PDA recognition by T cells was only modestly improved when MHC-I expression was enhanced by pretreatment with IFNγ. The fact that Msln was produced by all PDA lines suggests that these lines develop defects in antigen processing, and perhaps specifically Msln_406-414_, and that defective antigen processing and presentation appears to be a conserved mechanism of immune evasion. Alternatively, Msln epitopes may undergo post-translational modification – a process that has been revealed in studies of other antigens (99–103). Notably, even in conditions where target cell recognition was optimized, T cells failed to produce the essential effector cytokine, IFNγ. This suppressive phenomenon was largely independent of TGFβ and was consistently observed in diverse PDA lines from *KPC* tumors, perhaps suggesting that the suppression is driven by a mutant *KRAS*-and/or *TP53*-driven mechanism. Future studies aimed at identifying the presumably tumor intrinsic suppressive mechanisms are underway and have the potential to reveal a conserved immunotherapeutic target in PDA.

Based on findings that Msln antigen presentation restricted ACT efficacy, we generated a *KPC* cell line that overexpressed the cognate Msln_406-414_ epitope recognized by the 1045 TCR. As predicted, this strategy improved T cell recognition of tumor cells *in vitro* and *in vivo*. When combined with ACT using T cells lacking TGFβ signaling, mice exhibited improved survival. Aligning with prior models, *Tgfbr2*^-/-^ T cells were 5-fold more prevalent within tumors, adopted markedly increased effector and exhausted phenotypes, and were 4-fold more functional upon restimulation. Nevertheless, PDA tumors progressed, and mice succumbed to disease, as the orthotopic and invasive *KPC* PDA models are particularly aggressive. Our findings on limited Msln antigen processing/presentation call for more nuanced screening techniques when evaluating patients for TCR-engineered ACT that extend beyond measurement of target protein expression and more specifically assess target peptide antigen presentation on MHC. Additionally, approaches to increase antigen processing and presentation may be required for optimal efficacy of TCR-engineered ACT. As the exact defect in Msln antigen processing remains undefined, efforts to design therapies to restore antigen presentation will first require identification of the mechanisms that limit this process. Should, for example, these mechanisms be related to epigenetic suppression of proteins involved in antigen processing, the ability for histone deacetylase or DNA methyltransferase inhibitors to restore antigen presentation could be evaluated and potentially combined with ACT.

Overall, our studies reveal that while TGFβ is a potent target for immunotherapy in PDA, loss of sensitivity to this suppressive cytokine in ACT T cells is insufficient to facilitate cure of this highly aggressive and resistant disease. Thus, while targeting TGFβ can overcome some of the limitations of ACT in PDA, additional strategies are needed to further enhance the efficacy of this still promising approach to a nearly universally deadly disease.

