## Supplemental Figures and Table for "Abrogating TGFβ signaling in TCR-engineered T cells and enhancing antigen processing by tumor cells promotes sustained therapeutic activity in pancreatic ductal adenocarcinoma"

**SUPPLEMENTAL FIGURE LEGENDS**

**Supplemental Figure 1. 1045 TRex *Tgfbr2* KO T cells differentiate into effectors in an antigen-specific manner. A-C)** Representative plots (A) depicting $CD4^+$ (B) or $CD8^+$ (C) frequency of $CD45^+$ cells within inguinal LNs or spleens from 1045 TRex WT or KO mice. **D)** $V\beta9^+$ frequency of $CD8^+$ T cells. **E-F)** Representative plots (E) of $CD44^-CD62L^+$ $T_{Naïve}$, $CD44^+CD62L^+$ $T_{CM}$, or $CD44^+CD62L^-$ $T_{EFF}$ frequencies within $V\beta9^-CD8^+$ or $V\beta9^+CD8^+$ cells (F). **G)** $Klrg1^+$ frequency of $V\beta9^-CD8^+$ or $V\beta9^+CD8^+$ cells. **H-I)** Representative plots (H) depicting $Cx3cr1^+$ frequency of $V\beta9^-CD8^+$ or $V\beta9^+CD8^+$ cells. Dots are individual animals and n=5/group in all panels with mean ± SEM shown. Student's t-test was used for comparisons in all panels.

**Supplemental Figure 2. TGFβ limits T cell expansion without influencing central memory development. A)** Schematic illustrating ACT with co-transfer of 1045 TRex WT and KO cells ($2.5 \times 10^6$ of each cell type) with BiVax in healthy WT mice. Peripheral blood was obtained on a weekly basis for six weeks following ACT and tissues were isolated eight weeks after ACT. **B)** Frequency of $Thy1.1^+/1.1^+$ 1045 TRex WT cells and $Thy1.1^+/1.2^+$ 1045 TRex KO cells prior to ACT. **C)** Representative plot of circulating donor and endogenous (endo) $CD8^+$ T cells on day 28. **D)** Circulating 1045 TRex WT or KO donor frequency of $CD8^+$ T cells. **E)** Number of donor cells per μL of blood. **F)** Ratio of circulating 1045 TRex KO to WT cells. **G-J)** Representative plots (G) of $CD127^+CD62L^+$ $T_{CM}$ (H), $CD127^+CD62L^-$ $T_{EM}$ (I), or $CD127^-CD62L^-$ $T_{EFF}$ (J) frequency of 1045 TRex WT or KO donor cells. Dots are individual animals and n=3/group in all panels with mean ± SEM shown. Student's t-test was used for comparisons in all panels.

**Supplemental Figure 3. Tissue resident memory formation is largely maintained in the absence of TGFβ. A)** Frequency of $Thy1.1^+/1.1^+$ 1045 TRex WT cells and $Thy1.1^+/1.2^+$ 1045 TRex KO cells within the extravascular (i.e., $IV^-$) $CD8^+$ compartment of inguinal LN, spleen, pancreas (Panc), lung, liver, and small intestine intraepithelial lymphocytes (SI-IEL) of mice depicted in Supplemental Figure 2A at day 56 (8 weeks). **B)** Number of 1045 TRex WT or KO cells per organ. **C)** $CD103^+$ frequency of 1045 TRex WT or KO donor cells. **D)** $CD69^+$ frequency of 1045 TRex WT or KO donor cells. **E)** $CD49a^+$ frequency of 1045 TRex WT or KO donor cells. **F)** $Cxcr6^+$ frequency of 1045 TRex WT or KO donor cells. **G)** $CD69^+CD49a^+$ frequency of 1045 TRex WT or KO donor cells. **H)** $CD49a^+Cxcr6^+$ frequency of 1045 TRex WT or KO donor cells. **I)** $CD69^+Cxcr6^+$ frequency of 1045 TRex WT or KO donor cells. **J)** $CD69^+CD49a^+Cxcr6^+$ frequency of 1045 TRex WT or KO donor cells. Representative plots are shown in the upper portions of C-I. Dots are individual animals and n=3/group in all panels with mean ± SEM shown. Student's t-test was used for comparisons between two groups in all panels and one-way ANOVA with Tukey posttest was used for comparisons that included SI-IEL in C-I.

**Supplemental Figure 4: KPC tumor lines express Msln, MHC-I, and TGFβ. A)** Threshold cycles (Ct) for *Msln* transcripts in *KPC*2, *KPC*271, and *KPC*451 PDA tumor cell lines (left) or fold change relative to *KPC*2 (right). **B)** Membrane-bound (MB) and intracellular (IC) Msln protein in *KPC*2, *KPC*271, and *KPC*451 cells with secondary antibody only staining as a negative control. **C)** $H2D^b$ in unstimulated *KPC*2, *KPC*271, and *KPC*451 cells and *KPC* cells stimulated for 24 hours with IFNγ (100 μg/mL). MFI values are shown. **D)** Ct for *Tgfb1* in *KPC*2, *KPC*271, and *KPC*451 cells (left) or fold change relative to *KPC*2 (right). **E-F)** Representative images at 10X (E) or 40X (F) magnification from immunofluorescent microscopy of *KPC*2 tumors isolated from control mice on day 15 evaluating cytokeratin (CK), latency activated protein (LAP), and DAPI. **G)** Schematic showing generation of *KPC*2$^{Msln\text{-}ER}$, which is engineered to produce $Msln_{406\text{-}414}$ peptide targeted to the endoplasmic reticulum. A and D show mean ± SD from experimental triplicates and are representative of at least three independent experiments. B-C show representative findings from three independent experiments.

### Supplemental Figure 1

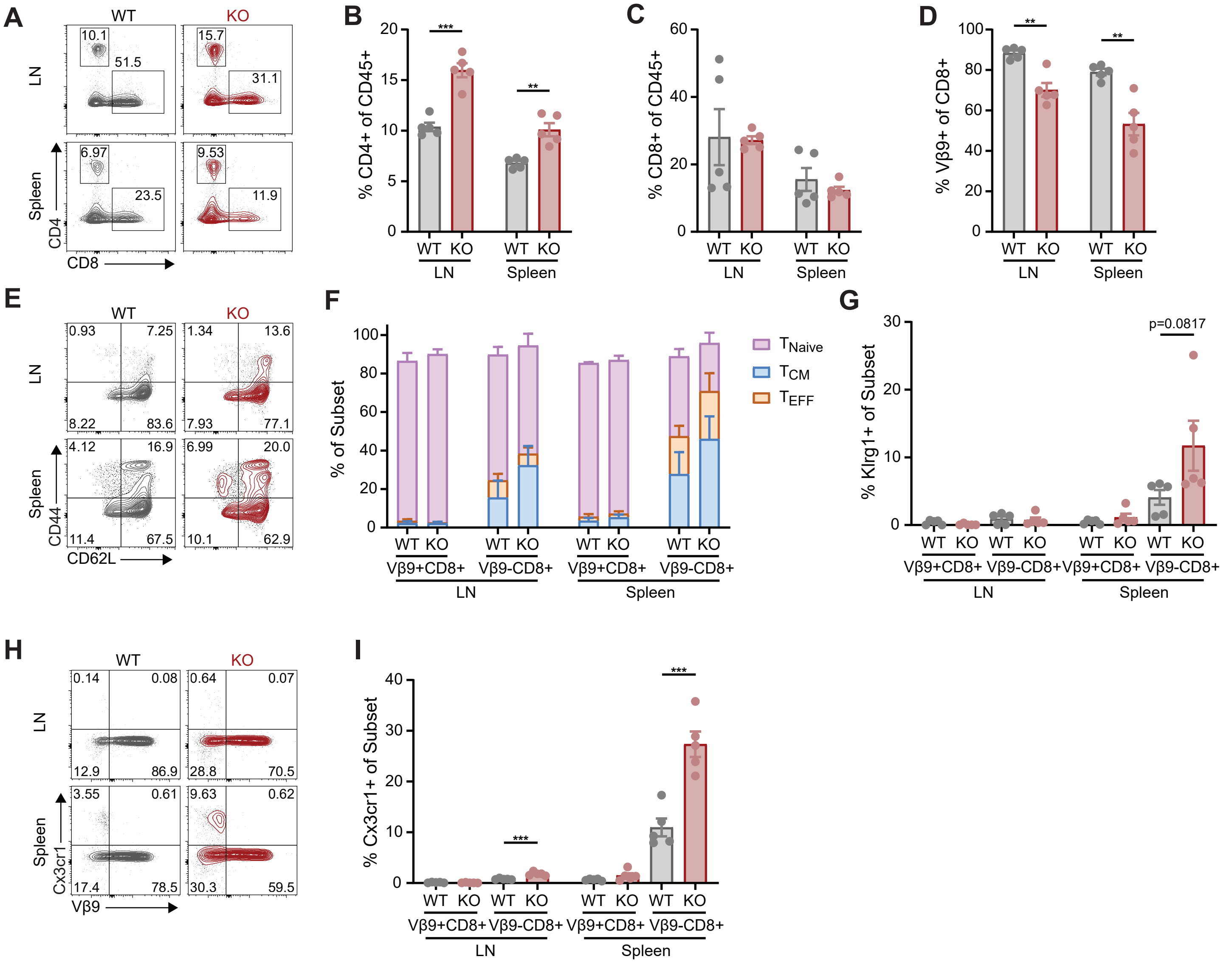

### Supplemental Figure 2

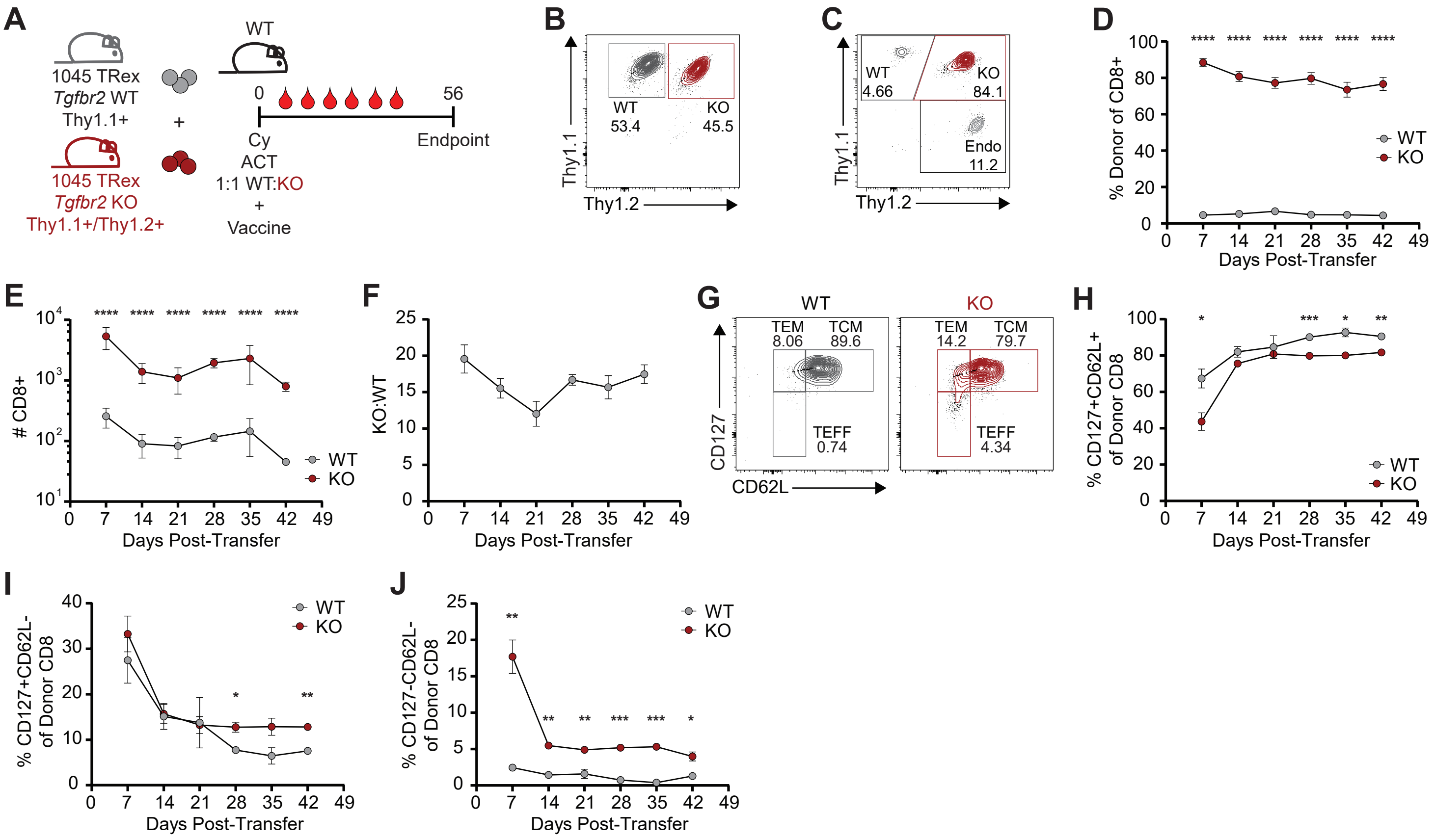

#### Supplemental Figure 3

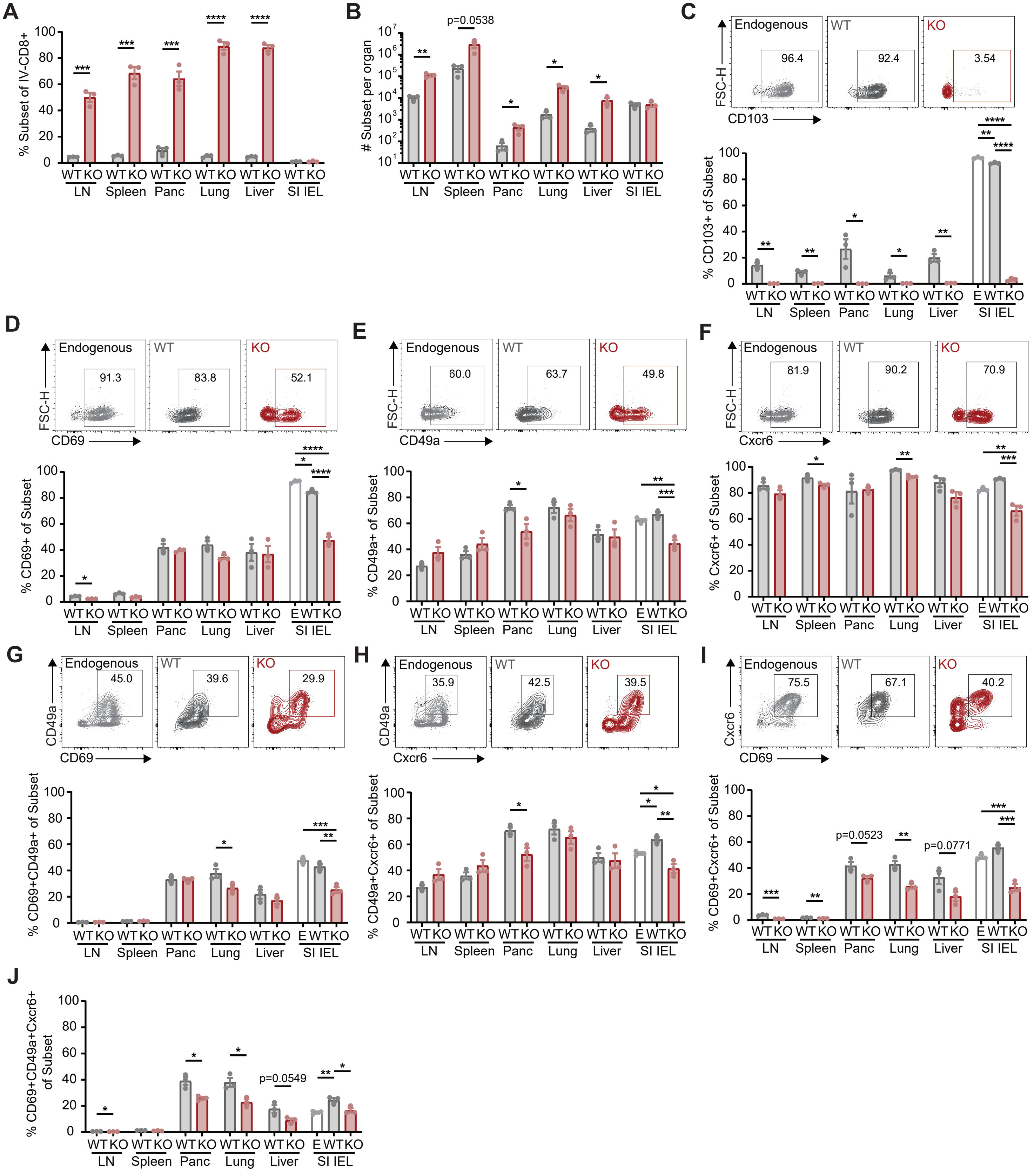

#### Supplemental Figure 4

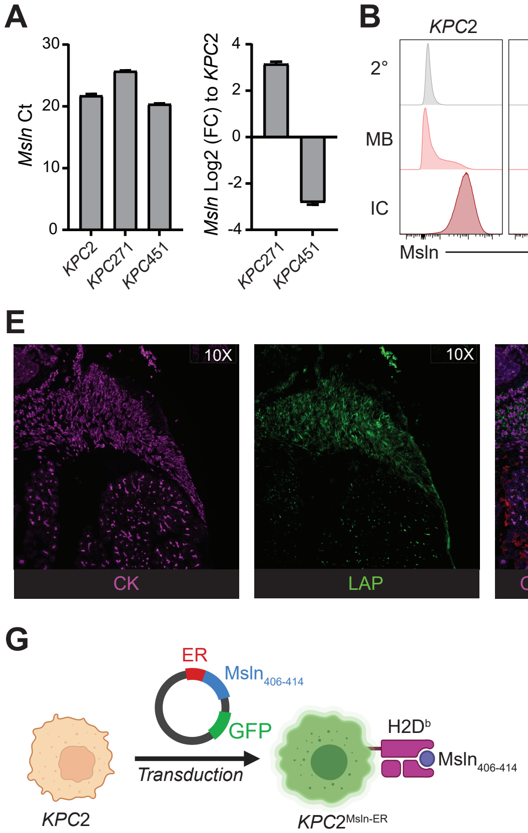

**Supplemental Table 1**

| Flow Cytometry and Immunofluorescent Microscopy Antibodies | | | | |
|---|---|---|---|---|
| **Fluorophore** | **Antibody Target** | **Clone** | **Vendor** | **Product #** |
| TNFa | BV711 | MP6-XT22 | BioLegend | 506349 |
| BUV563 | CD103 | 2 E7 | Invitrogen | 365-1031-82 |
| BV786 | CD103 | M290 | BD | 564322 |
| PE-Fire 640 | CD127 | S18006K | BioLegend | 158214 |
| BV421 | CD25 | PC61 | BioLegend | 102034 |
| BV650 | CD3 | 17A2 | BioLegend | 100229 |
| BUV496 | CD4 | GK1.5 | BD | 612952 |
| BV480 | CD4 | RM4-5 | BD | 565634 |
| BV785 | CD4 | GK1.5 | BioLegend | 100453 |
| PE-Cy5 | CD4 | GK1.5 | BioLegend | 100410 |
| Alexa Fluor 700 | CD44 | IM7 | BioLegend | 103026 |
| BV480 | CD44 | IM7 | BD | 566116 |
| APC | CD45 | 30-F11 | BioLegend | 103112 |
| APC Fire 810 | CD45 | 30-F11 | BioLegend | 103174 |
| BUV805 | CD45 | 30-F11 | Invitrogen | 368-0451-82 |
| RB780 | CD49a | Ha31/8 | BD | 755334 |
| BUV496 | CD62L | MEL-14 | Invitrogen | 364-0621-82 |
| BV605 | CD62L | MEL-14 | BioLegend | 104438 |
| BV650 | CD62L | MEL-14 | BioLegend | 104453 |
| BUV661 | CD69 | H1.2F3 | BD | 741478 |
| BUV737 | CD69 | H1.2F3 | BD | 612793 |
| PE-Dazzle 594 | CD69 | H1.2F3 | BioLegend | 104536 |
| APC | CD8a | 53-6.7 | BioLegend | 100712 |
| BUV395 | CD8a | 53-6.7 | BD | 563786 |
| BUV737 | CD8a | 53-6.7 | BD | 612759 |
| BV650 | CD8a | 53-6.7 | BioLegend | 100742 |
| NovaFluor Yellow 690 | CD8a | 53-6.7 | Invitrogen | M003T02Y05-A |
| BV570 | Cx3cr1 | SA011F11 | BioLegend | 149061 |

| | | | | |
|---|---|---|---|---|
| FITC | Cx3cr1 | SA011F11 | BioLegend | 149020 |
| PE-Cy7 | Cxcr3 | CXCR3-173 | BioLegend | 126516 |
| PerCP-Cy5.5 | Cxcr3 | CXCR3-173 | BioLegend | 126514 |
| FITC | Cxcr6 | SA051D1 | BioLegend | 151108 |
| PE-Dazzle 594 | Cxcr6 | SA051D1 | BioLegend | 151117 |
| Alexa Fluor 488 | Cytokeratin | C-11 | BioLegend | 628608 |
| FITC | Granzyme B | NGZB | Invitrogen | 11-8898-82 |
| PE-Cy7 | IFNg | XMG1.2 | Invitrogen | 25-7311-82 |
| BUV395 | Klrg1 | 2F1 | BD | 740279 |
| BV711 | Klrg1 | 2F1 | BioLegend | 138427 |
| Super Bright 780 | Klrg1 | 2F1 | Invitrogen | 78-5893-82 |
| BUV737 | Lag3 | C9B7W | BD | 741820 |
| BV650 | Lag3 | C9B7W | BioLegend | 125227 |
| BV510 | Live Dead | N/A | Tonbo | 13-0870 |
| Red 780 | Live Dead | N/A | Tonbo | 13-0865 |
| BUV395 | PD-1 | J43 | BD | 744549 |
| PE-Cy7 | PD-1 | 29F.1A12 | BioLegend | 135216 |
| BB700 | Slamf6 | 13G3 | BD | 742272 |
| BUV496 | Slamf6 | 13G3 | BD | 750046 |
| PE | Tcf1 | S33-966 | BD | 564217 |
| BV421 | Thy1.1 (CD90.1) | OX-7 | BioLegend | 202529 |
| PE | Thy1.1 (CD90.1) | OX-7 | BioLegend | 202524 |
| BUV737 | Thy1.2 (CD90.2) | 53-2.1 | BD | 741701 |
| BUV395 | Vb9 | MR10-2 | BD | 745620 |
| BV711 | Vb9 | MR10-2 | BD | 745452 |
| PE | Vb9 | MR10-2 | BioLegend | 139804 |

**Primers used for qPCR**

| **Target** | **Forward Sequence** | **Reverse Sequence** | **Assay ID** | **Vendor** |
|---|---|---|---|---|
| *Tgfb1* | TGATACGCCTGAGTGGCTGTCT | CACAAGAGCAGTGAGCGCTGAA | N/A | IDT |
| *Msln* | AGAGCCAGGAAAAGGCAGTCAG | CGATGGACTCATCCAACACTGC | N/A | IDT |
| *Gapdh* | N/A | N/A | Mm.PT.39a.1 | IDT |
